# Function of the HYDROXYCINNAMOYL-CoA:SHIKIMATE HYDROXYCINNAMOYL TRANSFERASE is evolutionarily conserved in embryophytes

**DOI:** 10.1101/2020.09.16.300285

**Authors:** Lucie Kriegshauser, Samuel Knosp, Etienne Grienenberger, Kanade Tatsumi, Desirée D. Gütle, Iben Sørensen, Laurence Herrgott, Julie Zumsteg, Jocelyn K.C. Rose, Ralf Reski, Danièle Werck-Reichhart, Hugues Renault

## Abstract

The plant phenylpropanoid pathway generates a major class of specialized metabolites and precursors of essential extracellular polymers that initially appeared upon plant terrestrialization. Despite its evolutionary significance, little is known about the complexity and function of this major metabolic pathway in extant bryophytes, which represent the non-vascular stage of embryophyte evolution. Here, we report that the *HYDROXYCINNAMOYL-CoA:SHIKIMATE HYDROXYCINNAMOYL TRANSFERASE* (*HCT*) gene that plays a critical function in the phenylpropanoid pathway during seed plant development, is functionally conserved in *Physcomitrium patens* (Physcomitrella), in the moss lineage of bryophytes. Phylogenetic analysis indicates that *bonafide* HCT function emerged in the progenitor of embryophytes. *In vitro* enzyme assays, moss phenolic pathway reconstitution in yeast and *in planta* gene inactivation coupled to targeted metabolic profiling, collectively indicate that *P. patens* HCT (PpHCT), similar to tracheophyte HCT orthologs, uses shikimate as a native acyl acceptor to produce a *p*-coumaroyl-5-*O*-shikimate intermediate. Phenotypic and metabolic analyses of loss-of-function mutants show that PpHCT is necessary for the production of caffeate derivatives, including previously reported caffeoyl-threonate esters, and for the formation of an intact cuticle. Deep conservation of HCT function in embryophytes is further suggested by the ability of *HCT* genes from *P. patens* and the liverwort *Marchantia polymorpha* to complement an *Arabidopsis thaliana* CRISPR/Cas9 *hct* mutant, and by the presence of phenolic esters of shikimate in representative species of the three bryophyte lineages.

## INTRODUCTION

Land colonization by plants, about 500 million years ago (Wickett et al., 2014; Puttick et al., 2018; Morris et al., 2018), was one of the most important evolutionary events associated with terraformation. Through photosynthetic activity and rock weathering, early land plants contributed to the rise of atmospheric oxygen, carbon sequestration and the development of soils (Lenton et al., 2016; Porada et al., 2016; Retallack, 1997). Plant settlement on land therefore paved the way for the development of rich terrestrial ecosystems and the emergence of new life forms (Kenrick and Crane, 1997).

This transition from water to land exposed plants to challenging terrestrial conditions, such as drought, harmful levels of solar (UV) radiation, lack of buoyancy, extended temperature range, and novel pathogenic microorganisms (Rensing et al., 2008; de Vries and Archibald, 2018). Successful land colonization thus required specific developmental and metabolic adaptations (Reski, 2018). The formation of extracellular, or apoplastic, protective barriers was probably one of the most critical innovations of land plants, as they shield cells from damaging environmental insults and allow the formation of specialized structures required for water and nutrient management (e.g. cuticles and vasculature). In angiosperms, such structures are essentially comprised of four canonical hydrophobic biopolymers – cutin, suberin, sporopollenin and lignin – that reinforce and waterproof the polysaccharide-based cell wall (Nawrath et al., 2013).

Some precursors of these polymers are generated through the phenylpropanoid pathway, one of the most important branches of so-called plant specialized metabolism, which also allows the accumulation of powerful UV screens and antioxidants (Renault et al., 2019; Vogt, 2010; Weng and Chapple, 2010). The ability to synthesize phenylpropanoids evolved during the course of terrestrialization and is often regarded as a key adaptation by plants to life on land (Weng and Chapple, 2010; de Vries et al., 2017; Renault et al., 2019). The most common products generated by the phenylpropanoid pathway – flavonoids, soluble phenolic esters and biopolymer precursors – all derive from *p*-coumaroyl-CoA (**Fig. 1A**). This hub molecule is produced through the activities of three essential enzymes in the initial steps of the phenylpropanoid pathway: phenylalanine ammonia-lyase (PAL); cinnamate 4-hydroxylase (C4H), which belongs to cytochrome P450 family 73 (CYP73); and 4-coumarate:CoA ligase (4CL) (**Fig. 1A**). In flowering plants, further functionalization of the phenolic ring requires shikimate ester intermediates and a two-enzyme module involving hydroxycinnamoyl-CoA:shikimate hydroxycinnamoyl transferase (HCT), which catalyzes transfer of the *p*-coumaroyl moiety from *p*-coumaroyl-CoA to shikimate (Hoffmann et al., 2003, 2004), and a second cytochrome P450, *p*-coumaroyl-shikimate 3’-hydroxylase (C3’H or CYP98), to generate caffeoyl-shikimate (Schoch et al., 2001; Franke et al., 2002; Alber et al., 2019) (**Fig. 1A**). HCT was shown to transfer the caffeoyl moiety back to coenzyme A to form caffeoyl-CoA *in vitro* (Hoffmann et al., 2003; Vanholme et al., 2013). Later studies reported the existence of a second route towards caffeoyl-CoA in flowering plants, involving the combined action of caffeoyl shikimate esterase (CSE) and 4CL (Vanholme et al., 2013; Saleme et al., 2017; Ha et al., 2016) (**Fig. 1A**). Recently, a bifunctional *p*-coumarate 3-hydroxylase/ ascorbate peroxidase (C3H/APX) was also characterized, revealing a metabolic shunt for phenolic ring 3-hydroxylation, directly from free *p*-coumaric acid (Barros et al., 2019) (**Fig. 1A**).

**Figure 1.**
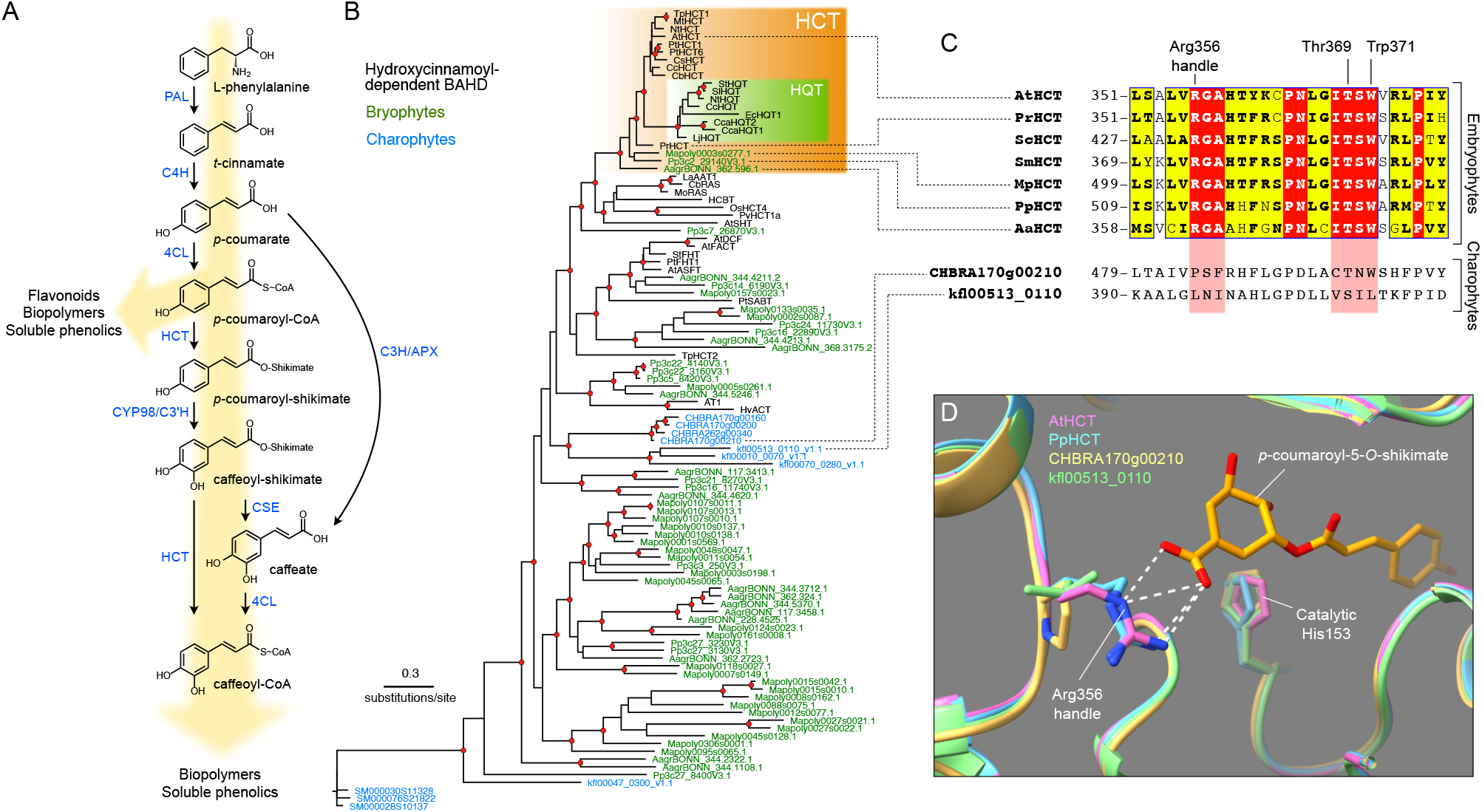
Evolutionary history of the HCT gene family. (A) Schematic representation of the phenylpropanoid pathway of angiosperms leading to caffeoyl-CoA. Enzyme names are indicated in blue. PAL, phenylalanine ammonia lyase; C4H, cinnamate 4-hydroxylase; 4CL, 4-coumarate:CoA ligase; HCT, hydroxycinnamoyl-CoA:shikimate hydroxycinnamoyl transferase; C3’H, *p*-coumaroyl ester 3’hydroxylase; CSE, caffeoyl-shikimate esterase; C3H/APX, *p*-coumarate 3-hydroxylase/ascorbate peroxidase. (B) Unrooted protein tree describing the phylogenetic relationships between 34 hydroxycinnamoyl-CoA-dependent BAHD acyltransferases of known biochemical function and all BAHDs from *Klebsormidium nitens, Chara braunii, Spirogloea muscicola, P. patens, Marchantia polymorpha and Anthoceros agrestis*. The tree is drawn to scale. Red dots indicate a maximum-likelihood ratio test branch support ≧ 0.80. Lists of characterized and uncharacterized BAHD proteins are available in Supplemental Tables S1 and S2, respectively. (C) Multiple sequence alignment highlighting the region comprising the three residues controlling shikimate acylation in selected HCT orthologs from the five major embryophyte groups. Corresponding regions from charophyte HCT homologs are displayed at the bottom for comparison. ScHCT, *Salvinia cucullata Sacu_v1.1_s0010.g004618*; SmHCT, *Selaginella moellendorfii 152997*. (D) Overlay of protein three-dimensional structures depicting the Arg356 handle interaction with shikimate. Models of *P. patens* (PpHCT, *Pp3c2_29140*, blue), *C. braunii* (*CHBRA170g00210*, yellow) and *K. nitens* (*kf/00513_0110*, green) HCT homologs were reconstructed using the crystal structure of AtHCT in complex with *p*-coumaroyl-5-*O*-shikimate (PDB entry: 5kju; pink). Residues are numbered according to AtHCT. White dashed lines represent predicted hydrogen bonds.

HCT belongs to clade V of the BAHD acyltransferase superfamily, which features enzymes that use coenzyme A-activated acyl donors and chemically diverse acceptors, such as organic acids, amines or fatty acids (D’Auria, 2006). Clade V also includes the closely-related enzymes hydroxycinnamoyl-CoA:quinate hydroxycinnamoyl transferases (HQT), which use quinate as a preferred acceptor, rather than shikimate. HQT is involved in the production of chlorogenic acids (caffeoyl-quinates), which are widespread among angiosperms, though absent from *Arabidopsis thaliana* (Niggeweg et al., 2004; Guo et al., 2014). Unlike caffeoyl-shikimate, caffeoyl-quinate is not considered to be a key intermediate in lignin biosynthesis, but rather involved in responses to biotic and abiotic stressors, especially UV radiation (Clé et al., 2008; Niggeweg et al., 2004). An investigation of HCT catalytic properties revealed broad acceptor substrate permissiveness, extending beyond shikimate (Hoffmann et al., 2004, 2003; Sander and Petersen, 2011; Eudes et al., 2016). However, structural studies of HCT/HQT uncovered key amino acid residues that control shikimate and/or quinate acylation, thereby specifying the two types of enzymes (Levsh et al., 2016; Lallemand et al., 2012; Chiang et al., 2018). HCT represents a pivotal step in controlling lignin biosynthesis and composition, as demonstrated by *HCT* silencing studies in seed plants that consistently alter lignin content and/or composition, and often lead to adverse effects on growth (Hoffmann et al., 2004; Wagner et al., 2007; Besseau et al., 2007; Gallego-Giraldo et al., 2011).

Bryophytes and charophyte algae, the embryophyte sister group, are devoid of lignin, although some of them seem to possess parts of the genetic toolkit required to synthesize phenolic intermediates (de Vries et al., 2017; Renault et al., 2019; Jiao et al., 2020). The nature and the role of such early phenylpropanoid derivatives are still poorly documented. We recently showed through a molecular genetic approach, targeting the *Physcomitrium patens* (Physcomitrella) *CYP98* gene, that a moss phenylpropanoid pathway is involved in the synthesis of caffeate precursors necessary to support cuticular biopolymer formation and erect (3D) growth (Renault et al., 2017). The major acylated products formed by the moss were shown to be threonate esters (*p*-coumaroyl-threonate and caffeoyl-threonate), while shikimate and quinate esters were not detected (Renault et al., 2017). However, a survey of embryophyte CYP98 substrate preference *in vitro* showed that the moss enzyme poorly converts *p*-coumaroyl-threonates, compared with *p*-coumaroyl-shikimate (Alber et al., 2019), leaving the nature of the native pathway intermediates in the moss unclear. Here, we sought to address this question by performing a functional analysis of a candidate *P. patens HCT* gene, which encodes the enzyme generating the CYP98 substrate. Combining *in silico*, *in vitro* and *in vivo* analyses, we demonstrate conservation of HCT catalytic properties and physiological function across the 500 million years of embryophyte evolution.

## RESULTS

### *A bona fide HCT* gene emerged in an embryophyte progenitor and was subsequently conserved

We performed a search for potential *HCT* genes in fully sequenced charophyte and bryophyte genomes. All BAHD acyltransferase protein sequences from the charophytes *Klebsormidium nitens* (*Klebsormidiophyceae*), *Chara braunii* (*Characeae*) and *Spirogloea muscicola*, (*Zygnematophyceae*) and from the bryophytes *P. patens* (moss), *Marchantia polymorpha* (liverwort) and *Anthoceros agrestis* (hornwort) were aligned with 34 already characterized hydroxycinnamoyl-CoA-dependent BAHD transferases prior to phylogeny reconstruction (complete lists in **Tab. S1 and S2**). The resulting protein tree structure revealed a well-supported HCT clade with single members for each bryophyte species at its root (**Fig. 1B**). The angiosperm-specific HQT proteins clustered as a sister group to angiosperm HCTs, suggesting they originated from *HCT* duplication (**Fig. 1B**). BAHD enzymes from the charophytes *C. braunii* and *K. nitens* were not closely associated with HCTs, but rather occupied a basal position with respect to characterized hydroxycinnamoyl-CoA-dependent BAHD. Proteins from the Zygnematophyceae *S. muscicola* were found to be even more divergent from characterized HCT proteins (**Fig. 1B**). The multiple protein alignment revealed a strict conservation in representative embryophyte HCTs of the three residues (Arg356, Thr369 and Trp371) previously shown to be critical for HCT activity (Lallemand et al., 2012; Levsh et al., 2016; Chiang et al., 2018), whereas their conservation was only partial in charophyte homologs (**Fig. 1C, Fig. S1**). More particularly, the Arg356 handle was absent from charophyte BAHDs (**Fig. 1C, Fig. S1**). Finer details were gained through homology-modelling of HCT candidate proteins from *P. patens* (PpHCT, *Pp3c2_29140*), *C. braunii* (*CHBRA170g00210*) and *K. nitens* (*kfl00513_0110*) using the crystal structure of *Arabidopsis thaliana* AtHCT in complex with *p*-coumaroyl-5-*O*-shikimate as a template. The predicted protein structures indicated that, similar to AtHCT, PpHCT binds the shikimate carboxyl group through an arginine handle (**Fig. 1D**). In charophyte proteins, the critical arginine residue was replaced by proline (*CHBRA170g00210*) or leucine (*kfl00513_0110*), neither of which is predicted to form hydrogen bonds with shikimate (**Fig. 1D**). Taken together, these data point to the emergence of *bona fide HCT* genes in the last common ancestor of embryophytes about 500 Ma, concurrently with the appearance of cuticles (Philippe et al., 2020) and prior to the capacity to produce the phenolic biopolymer lignin.

### *PpHCT* is co-expressed with the *p-coumaroyl ester 3’-hydroxylase PpCYP98*

We then undertook a functional study of the *P. patens HCT* candidate gene (*PpHCT, Pp3c2_29140*) identified in the phylogenetic analysis. First, we used publicly available co-expression data derived from the *P. patens* gene atlas project (Perroud et al., 2018). This indicated that the expression profile of *PpHCT* is tightly correlated with those of two genes encoding enzymes flanking the HCT step, potentially forming a core enzymatic module in the moss phenolic pathway: *4-coumarate-CoA ligase 1* (*Pp4CL1*; *Pp3c18_6360;* Silber et al., 2008) and the functionally characterized *PpCYP98* (*Pp3c22_19010*; Renault et al., 2017) (**Fig. 1A, 2A**). These data were supported by our qRT-PCR analysis, which showed higher (at least 4-fold) expression levels of all three genes in gametophores than in protonema (**Fig. 2B**). To increase the spatial resolution of the *PpHCT* expression analyses, we generated knock-in *PpHCT:uidA* reporter lines by homologous recombination (**Fig. S2**). GUS staining was found to be restricted to the apex of both young and old gametophores (**Fig. 2C**), which is very similar to the previously reported *PpCYP98:uidA* GUS staining profile (Renault et al., 2017).

**Figure 2.**
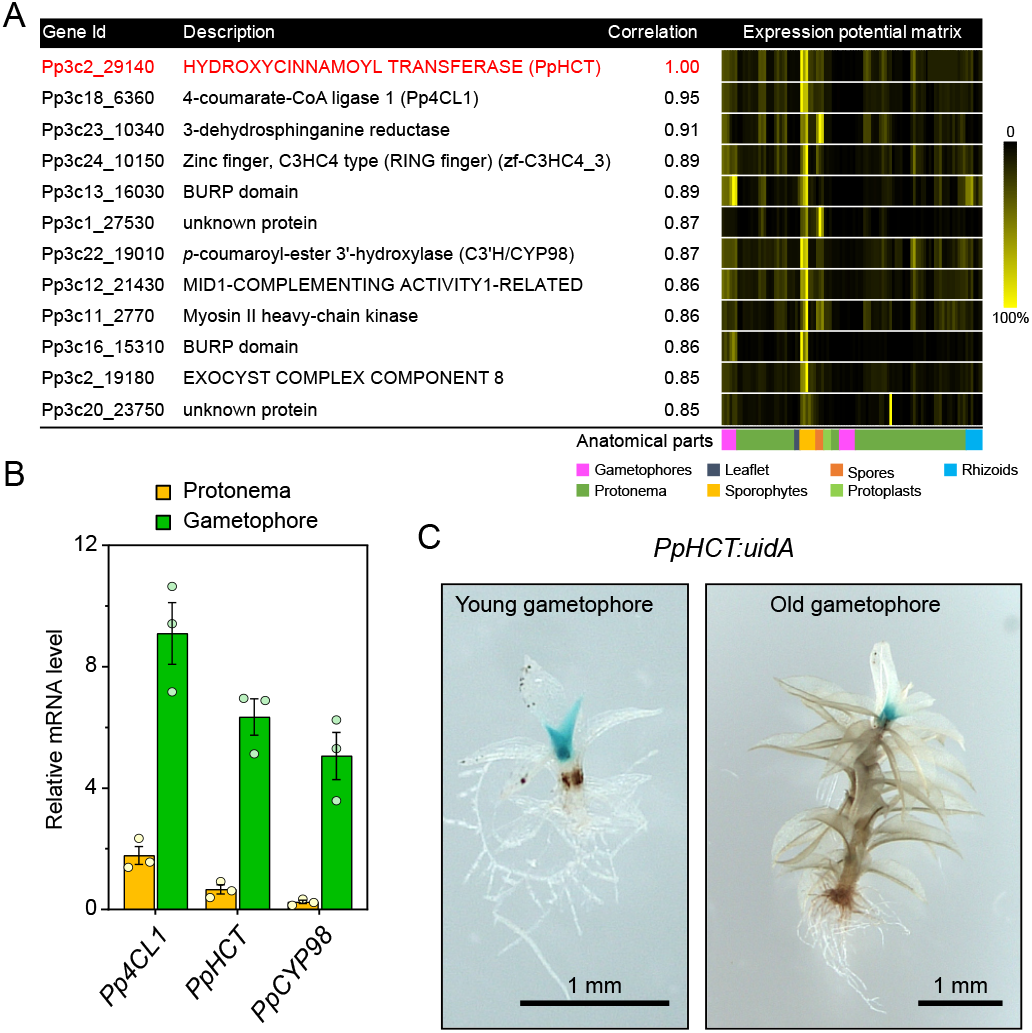
*PpHCT* co-expression network and expression pattern. (A) List of genes co-expressed with *PpHCT* (*Pp3c2_29140*). Each element of the matrix represents the expression potential (% of maximum expression across the matrix) of a given gene (line) in a defined condition (column) derived from various anatomical parts. Data are retrieved from the *P. patens* gene atlas project (Perroud et al., 2018). (B) qRT-PCR monitoring of *Pp4CL1*, *PpHCT* and *PpCYP98* expression in protonema and gametophore tissues. Data are the mean ± SEM of three biological replicates derived from three independent cultures. (C) GUS staining pattern in *PpHCT:uidA* lines indicating a prominent expression in the apex of gametophores.

### PpHCT demonstrates substrate permissiveness *in vitro*

Previous data suggested that phenolic esters of threonic acid are the most likely intermediates in the *P. patens* phenylpropanoid pathway (Renault et al., 2017). Accordingly, we hypothesized that PpHCT generates *p*-coumaroyl-threonate from *p*-coumaroyl-CoA and L-threonic acid, and we tested this with *in vitro* assays using recombinant PpHCT expressed in *Escherichia coli*. We observed that PpHCT produced mainly *p*-coumaroyl-4-*O*-threonate and a minor amount of *p*-coumaroyl-2-*O*-threonate *in vitro*, indicative of substantial regiospecificity (**Fig. 3A-B**). We then tested shikimic acid and quinic acids as acyl acceptors, since they are native and accepted substrates of tracheophyte HCTs, respectively (Hoffmann et al., 2003; Chiang et al., 2018). PpHCT catalyzed the transfer of *p*-coumarate from *p*-coumaroyl-CoA to both of them (**Fig. 3A**). A strong regiospecificity favored the 5-position for shikimate and quinate acylation (**Fig. 3A-B**). We then investigated PpHCT acyl-CoA donor preference using end-point enzyme assays, testing all pairwise combinations of the donors *p*-coumaroyl-CoA, caffeoyl-CoA and feruloyl-CoA with the acceptors threonate, shikimate and quinate. As shown in **Figure 3C**, PpHCT used all three acyl donors when shikimate was the acceptor, and the highest activity was observed with the combination of shikimate and feruloyl-CoA. PpHCT thus displayed significant donor and acceptor permissiveness. This was more pronounced with *p*-coumaroyl-CoA, the *P. patens* native acyl donor, which in addition was the only donor that coupled with threonate (**Fig. 3C**). A striking difference between PpHCT and orthologs from vascular plants is the presence of a 144-amino acid flexible loop joining the two main folded domains of the protein (**Fig. S3**). To test the impact of this loop on enzyme activity, a truncated version of PpHCT lacking the internal loop domain was produced in *E. coli* (**Fig. S4**). The truncation did not affect substrate preference; however, it caused a minor decrease in activity with threonate as the acceptor, without altering the shikimate and quinate acylation activity (**Fig. S4**).

**Figure 3.**
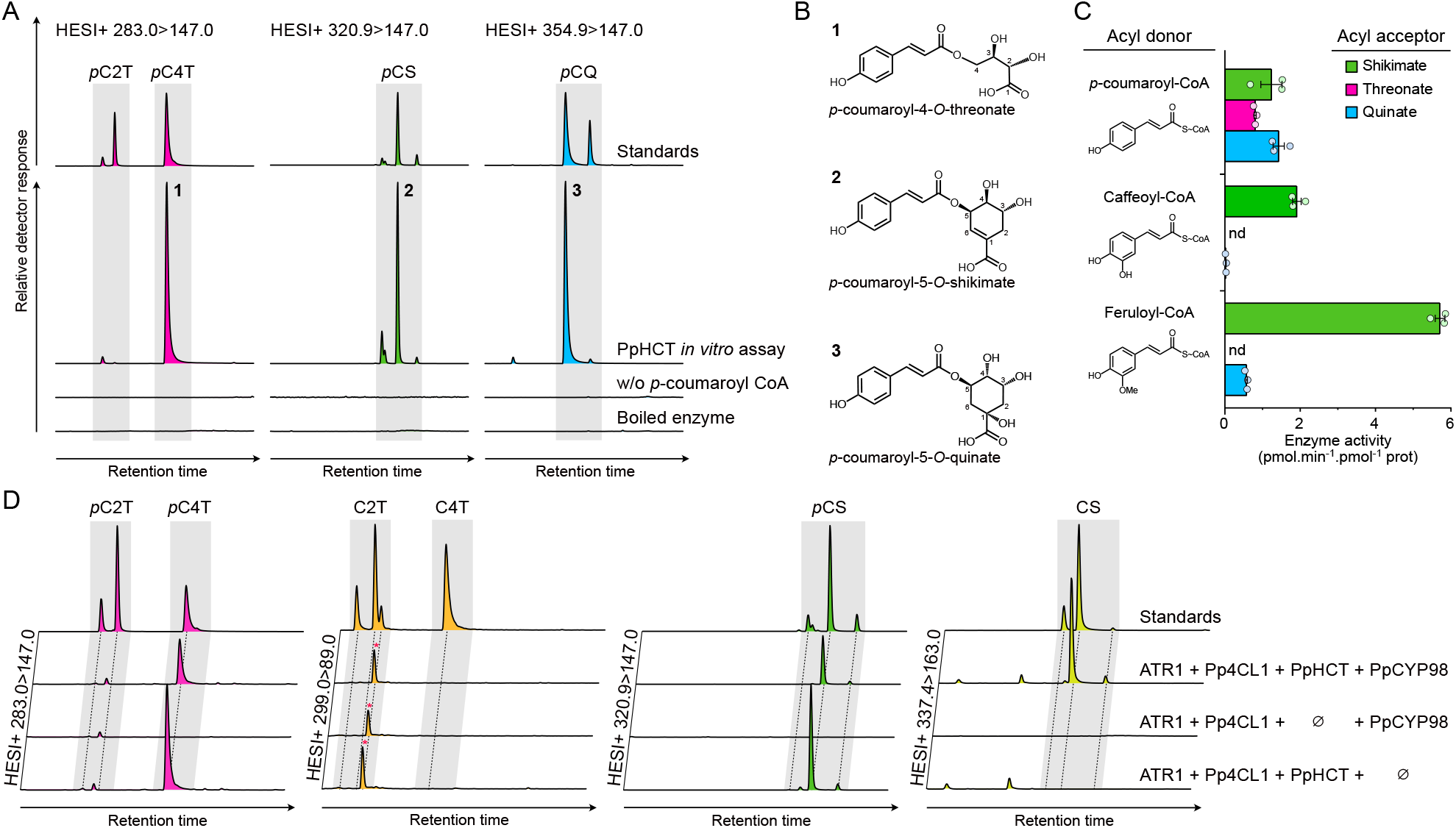
Investigation of recombinant PpHCT catalytic properties. (A) Overlay of UHPLC-MS/MS chromatograms showing the production of *p*-coumaroyl esters by PpHCT *in vitro.* The recombinant protein was incubated with *p*-coumaroyl-CoA and one of the three different acyl acceptors: quinate, shikimate and threonate. Assays with boiled enzyme or without *p*-coumaroyl-CoA were used as controls. *p*C2T, *p*-coumaroyl-2-*O*-threonate; *p*C4T, *p*-coumaroyl-4-*O*-threonate; *p*CS, *p*-coumaroyl-shikimate; *p*CQ, *p*-coumaroyl-quinate. (B) Structures of the main *p*-coumaroyl esters detected in (A). (C) PpHCT activity for all pairwise combinations involving *p*-coumaroyl-CoA, caffeoyl-CoA and feruloyl-CoA as acyl donor, together with quinate, threonate and shikimate as an acyl acceptor. Enzyme activity was calculated based on end-point assays analyzed by HPLC-UV. Data are the mean ± SEM of three independent enzyme assays. nd, not detected. (D) Overlay of UHPLC-MS/MS chromatograms showing the production of phenolic esters in whole-cell assays using engineered *Saccharomyces cerevisae* strains expressing different combinations of *Pp4CL1*, *PpHCT*, *PpCYP98* and *ATR1* genes. *p*-coumaroyl-threonates (*p*C2T, *p*C4T), caffeoyl-threonates (C2T, C4T), *p*-coumaroyl-shikimates (*p*CS) and caffeoyl-shikimates (CS) esters were simultaneously analyzed from yeast culture supplemented with *p*-coumarate and L-threonate using HESI+ MRM methods. Y axes of yeast extract chromatographs are linked to show each phenolic ester. For caffeoyl-threonate, a non-specific signal was detected regardless of the gene set (red asterisks).

### PpHCT kinetic parameters largely favor shikimate acylation

To gain deeper insights into PpHCT catalytic properties, enzyme kinetic parameters were determined from activity saturation curves and Michaelis-Menten nonlinear regression (**Fig. S5**). We focused the kinetic analysis on the three acyl acceptors, threonate, quinate and shikimate, and on the native acyl donor *p*-coumaroyl-CoA. The results, summarized in **Table 1,** revealed an obvious preference of PpHCT for shikimate as an acyl acceptor, in terms of affinity (*K*_m_: 0.22 mM) and velocity (*k*_cat_: 5.1 s^−1^), compared with threonate (*K*_m_: 17.2 mM, *k*_cat_: 0.16 s^−1^). The calculated catalytic efficiency with shikimate (*k*_cat_/*K*_m_) was ~2,500-fold higher than with threonate (**Tab. 1**). PpHCT activity with quinate was mixed, exhibiting low affinity (*K*_m_: 9.4 mM) but a rather high velocity (*k*_cat_: 3.5 s^−1^). PpHCT affinity for *p*-coumaroyl-CoA was 60 μM when shikimate was used as acceptor, a value 7-times lower than when threonate was used as an acceptor (**Tab. 1**). PpHCT kinetic parameters with shikimate were found overall to be similar to those of angiosperm orthologs (Hoffmann et al., 2003; Levsh et al., 2016).

**Table 1.** Summary of PpHCT kinetic parameters. Enzyme affinity (*K*_m_) and velocity (*k*_cat_) constants were determined from PpHCT activity saturation curves, based on nonlinear Michaelis-Menten regression (**see Methods and Fig. S5**). Results are the means of three independent enzyme reactions; 95% confidence intervals (profile likelihood) are provided within brackets.

| Substrates | | $K_m$ (mM) | $k_{cat}$ (s<sup>-1</sup>) | $k_{cat}/K_m$ (s<sup>-1</sup>.M<sup>-1</sup>) |
| --- | --- | --- | --- | --- |
| Fixed | Variable |  |  |  |
| *p*-coumaroyl-CoA | Threonate | 17.2 (14.9-19.9) | 0.16 (0.15-0.17) | 9 |
|  | Shikimate | 0.22 (0.17-0.29) | 5.1 (4.8-5.4) | 23 005 |
|  | Quinate | 9.4 (6.7-13.7) | 3.5 (3.0-4.2) | 368 |
| Threonate | *p*-coumaroyl-CoA | 0.43 (0.27-0.76) | 0.18 (0.14-0.25) | 407 |
| Shikimate | *p*-coumaroyl-CoA | 0.06 (0.03-0.10) | 17.0 (14.1-20.7) | 283 333 |

The PpHCT kinetic parameters suggest that threonate would need to be present in much higher concentrations than shikimate in *P. patens* to be used as a substrate by PpHCT. To investigate this hypothesis, we first determined the absolute levels of the three potential acyl acceptors, (-)-shikimate, D-quinate and L-threonate, in the gametophyte of the three bryophytes *P. patens*, *Anthoceros agrestis* and *Marchantia polymorpha*, and in the angiosperm *Arabidopsis thaliana* by UHPLC-MS/MS (**see Methods section**). Levels of L-phenylalanine, the amino acid that initiates the phenylpropanoid pathway, and L-malate, an intermediate of the Krebs cycle, were simultaneously measured and served as metabolic benchmarks. Metabolic profiling revealed a higher level of threonate in *P. patens* than in the other species, while phenylalanine and malate levels remained similar among them (**Tab. S3**). At the level of the whole *P. patens* gametophore, threonate was found to be ~250-times more abundant than shikimate; in the other species, the threonate/shikimate ratio ranged from 0.1 to 2 (**Tab. S3**). Concentrations and ratios of these compounds might however differ in specific tissues, cell types or subcellular compartments. Quinate was not detected in any of the plant extracts, in accordance with the absence of a *bona fide* quinate dehydrogenase in the investigated species (Guo et al., 2014; Carrington et al., 2018). To test whether a 250-fold excess in threonate would shift the equilibrium from shikimate to threonate acylation, we incubated recombinant PpHCT with *p*-coumaroyl-CoA in the presence of 2.5 mM L-threonate and 10 μM (-)-shikimate, recapitulating a 250:1 molar ratio of competing substrates *in vitro*. Under these conditions, PpHCT activity was 37-times higher with shikimate than with threonate as an acyl acceptor (**Fig. S6A-B**). These results confirm that the catalytic properties of PpHCT favor shikimate acylation, even in presence of a far higher concentration of threonate.

### Reconstitution of the moss phenolic pathway in yeast

The *in vivo* functionality of PpHCT and its ability to operate with potential partner enzymes were investigated in engineered *Saccharomyces cerevisiae* co-expressing *Pp4CL1*, *PpHCT*, and *PpCYP98*, as well as *Arabidopsis thaliana ATR1* (*At4g24520*) encoding a P450 reductase to ensure sufficient electron supply to PpCYP98 (Urban et al., 1997). Since *S. cerevisiae* does not naturally synthesize phenylpropanoids or threonate, we supplemented the yeast culture media with *p*-coumarate and L-threonate 6h after the onset of galactose-induced recombinant protein production. UHPLC-MS/MS analysis of yeast culture extracts revealed the production of *p*-coumaroyl-4-*O*-threonate but not *p*-coumaroyl-2-*O*-threonate (**Fig. 3D**), consistent with PpHCT having a strong regiospecificity. Notably, caffeoyl-threonate was not detected in the yeast culture extracts (**Fig. 3D**). *S. cerevisiae* synthesizes shikimate as an intermediate of aromatic amino acid biosynthesis and, accordingly, we detected *p*-coumaroyl-shikimate in extracts of all *PpHCT*-expressing yeast strains (**Fig. 3D**), which confirmed PpHCT promiscuity *in vivo*. Caffeoyl-shikimate was readily detected in the yeast extracts, indicating that shikimate esters were intermediates, allowing an effective coupling of PpHCT and PpCYP98 activities (**Fig. 3D**). The major *p*-coumaroyl ester isomers produced in yeast were similar to those predominantly generated by PpHCT *in vitro* (**Fig. 3B**). In parallel, we used the yeast platform to assess the catalytic activity of the *K. nitens* HCT homolog kfl00513_0110 (see **Fig. 1**), and found that it did not lead to the production of detectable *p*-coumaroyl-shikimate when co-expressed with *Pp4CL1*, *PpCYP98* and *ATR1* (**Fig. S7**). This supports the idea that charophyte HCT homologous proteins do not act as canonical HCT enzymes.

### PpHCT efficiently converts caffeoyl-5-*O*-shikimate into caffeoyl-CoA *in vitro*

Earlier work on recombinant proteins from tobacco and Arabidopsis showed that *in vitro*, HCT catalyzes the conversion of caffeoyl-5-*O*-quinate or caffeoyl-5-*O*-shikimate into caffeoyl-CoA, using coenzyme A as acyl acceptor (Hoffmann et al., 2003; Vanholme et al., 2013); a reaction hereafter referred to as HCT reverse reaction (**Fig. 4A**). However, there has subsequently been little additional experimental support for the existence of this reaction, in particular in plant lineages other than angiosperms. We therefore tested *in vitro* the ability of recombinant PpHCT to catalyze the production of caffeoyl-CoA from caffeoyl esters. We first developed a sensitive HPLC-UV method that provides a linear response to caffeoyl-CoA (**Fig. 4B and Methods section**). We then performed *in vitro* assays with recombinant PpHCT, using coenzyme A as acyl acceptor and either caffeoyl-5-*O*-shikimate or caffeoyl-5-*O*-quinate as acyl donors. Caffeoyl-threonates, which are not commercially available and are not synthesized efficiently by CYP98 enzymes *in vitro* (Alber et al., 2019), could not be tested as substrates. HPLC-UV analysis of enzyme reaction products showed a sharp decrease of caffeoyl-5-*O*-shikimate in the PpHCT assay with a concomitant appearance of a peak that was identified as caffeoyl-CoA, based on retention time and absorbance spectrum comparison with an authentic standard (**Fig. 4C-D**). It was noted that minor amounts of additional caffeoyl-shikimate isomers were present in the reaction, but were not effectively metabolized by PpHCT (**Fig. 4C**). Based on caffeoyl-CoA production, PpHCT reverse activity with caffeoyl-5-*O*-shikimate as an acyl donor was comparable to that of the conjugating activity producing *p*-coumaroyl-5-*O*-shikimate. However, PpHCT reverse activity using caffeoyl-5-*O*-quinate as an acyl donor was seven-times lower (**Fig. 4E**), and this result was confirmed by the amount of caffeoyl esters remaining at the end of the enzyme assay (**Fig. 4F**).

**Figure 4.**
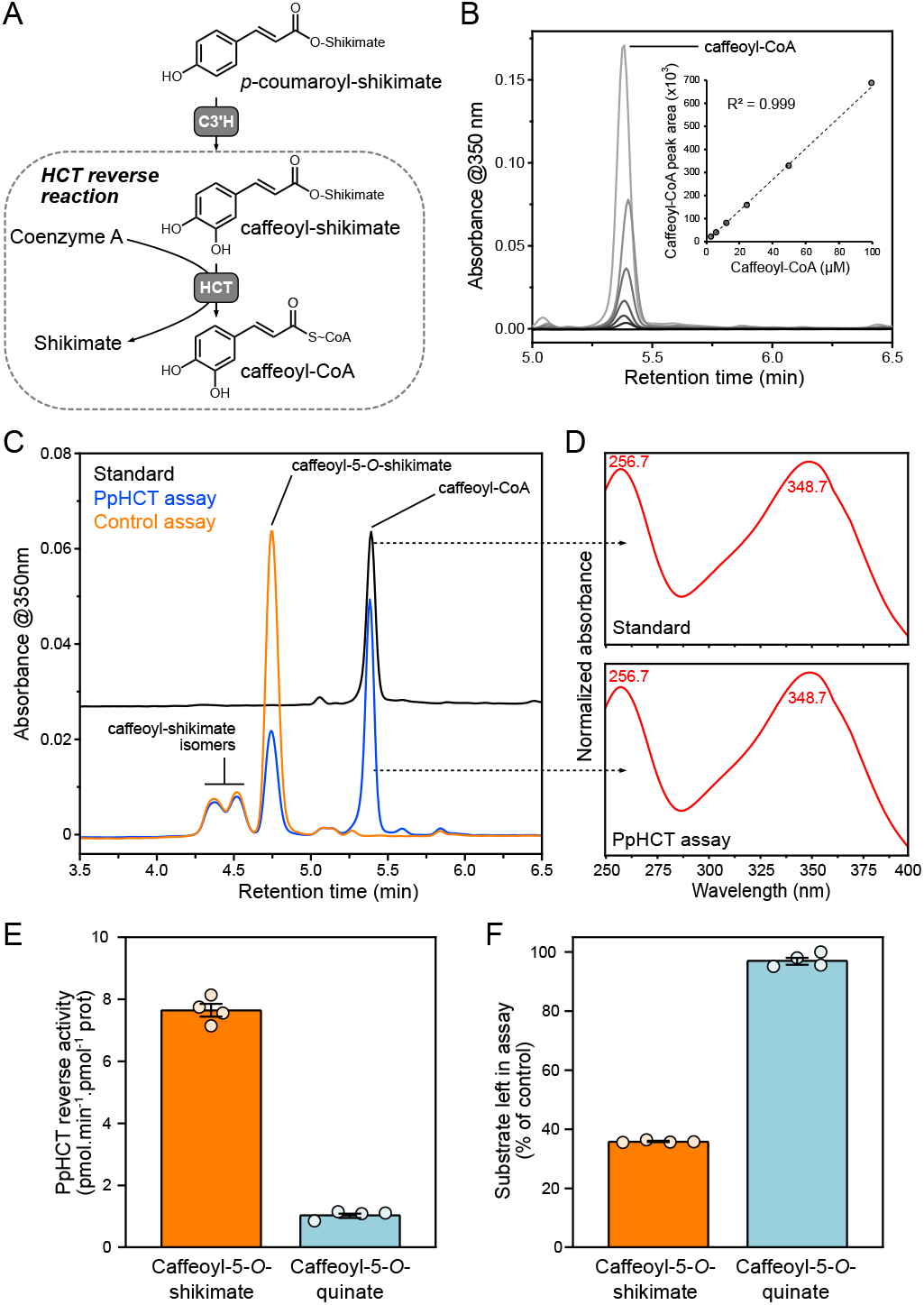
Investigation of PpHCT reverse reaction *in vitro*. (A) Scheme depicting the PpHCT reverse reaction that leads from caffeoyl-shikimate to caffeoyl-CoA thioester. (B) HPLC-UV detection of authentic caffeoyl-CoA molecule at 350 nm and associated calibration curve exhibiting linearity over at least three orders of concentration. (C) Representative HPLC-UV chromatograms showing the conversion of caffeoyl-5-*O*-shikimate into caffeoyl-CoA by PpHCT *in vitro*. Control assay was performed with boiled PpHCT enzyme. (D) Absorbance spectrum from 240 to 400 nm of PpHCT assay and caffeoyl-CoA standard peaks. Wavelengths corresponding to maximum absorbance are indicated. (E) PpHCT reverse activity with caffeoyl-5-*O*-shikimate or caffeoyl-5-*O*-quinate as acyl donor. (F) Relative level of caffeoyl-5-*O*-shikimate and caffeoyl-5-*O*-quinate left in PpHCT assay (% of control assay). Data are the mean ± SEM of four independent enzyme assays.

### PpHCT produces *p*-coumaroyl-shikimate *in planta* as a precursor of caffeate derivatives

Next, we generated four independent *PpHCT* deletion mutants (Δ*PpHCT*) via homologous recombination (**Fig. 5A**, **Fig. S8**) in order to address the *in planta* function of *PpHCT*. The four Δ*PpHCT* lines lacked full length *PpHCT* transcripts (**Fig. 5B**), leading to a complete abolishment of HCT activity in gametophore protein extracts (**Fig. 5C**). The Δ*PpHCT* lines phenocopied Δ*PpCYP98* mutants (Renault et al., 2017) and newly reported *PpHCT* loss-of-function mutant *Ppnog2-R* (Moody et al., 2020), characterized by defective gametophore development (**Fig. 5D-F**, **Fig. S8**). UV-fingerprinting of gametophore metabolite extracts revealed the absence of major peaks in Δ*PpHCT* mutant chromatogram (**Fig. 5G**). This low-resolution UV analysis was refined by targeted UHPLC-MS/MS analysis, which revealed both qualitative and quantitative differences in threonate esters. As expected, if PpHCT generates the substrate(s) of PpCYP98, caffeoyl-threonates were absent from Δ*PpHCT* (**Fig. 5H**). Unexpectedly, however, levels of *p*-coumaroyl-threonate esters were higher in the Δ*PpHCT* lines (**Fig. 5H**). Taken together, these data suggest that *p*-coumaroyl-threonate esters: (i) are not derived from PpHCT activity, implying the existence of another dedicated enzyme in *P. patens*; and (ii) are not the native substrates of PpCYP98, although they could be metabolized *in vitro* (Renault et al., 2017). We addressed the identity of this putative enzyme by testing the ability of each of the twelve full-length, expressed BAHD proteins from *P. patens* to catalyze the formation of *p*-coumaroyl-threonate in yeast. Only PpHCT was found to catalyze threonate acylation (**Fig. S9A**). This result was corroborated by *in vitro* assays with protein extracts from *P. patens* gametophores, which did not yield detectable *p*-coumaroyl-4-*O*-threonate (**Fig. S9B**). Next, to investigate the existence of potentially overlooked hydroxycinnamoyl intermediates, and in particular caffeoyl conjugates, gametophore extracts were submitted to acid hydrolysis to release hydroxycinnamate moieties prior to UHPLC-MS/MS analysis. Caffeate was not detected in mutant gametophore hydrolyzed extracts (**Fig. 5I**). This result indicates that PpHCT provides the only route toward caffeate derivatives in *P. patens*, in accordance with the absence of a *bona fide C3H/APX* gene in *P. patens* (Barros et al., 2019). A large increase in the amount of *p*-coumarate in hydrolyzed extracts (**Fig. 5I**) was consistent with the previously reported accumulation of *p*-coumaroyl-threonates in Δ*PpHCT* mutant lines (**Fig. 5H**). Taking advantage of the increased sensitivity and resolution provided by a new UHPLC-MS/MS analytical platform, we searched for shikimate esters in gametophore extracts in which we had not detected these compounds previously (Renault et al., 2017). To improve the detection threshold, extracts were also concentrated five-fold, and under these conditions we detected *p*-coumaroyl-5-*O*-shikimate in gametophore extracts from wild-type *P. patens*, but not in those from Δ*PpHCT* (**Fig. 5J-K**). The results were orthogonally validated by both retention time comparison with molecular standards and two different mass transitions in positive and negative modes (signal-to-noise ratio > 80). Taken together, the metabolic analysis of the Δ*PpHCT* mutants confirmed a key function of HCT in the production of caffeate derivatives in *P. patens* via the formation of a *p*-coumaroyl-5-*O*-shikimate intermediate and did not support the existence of alternative pathways.

**Figure 5.**
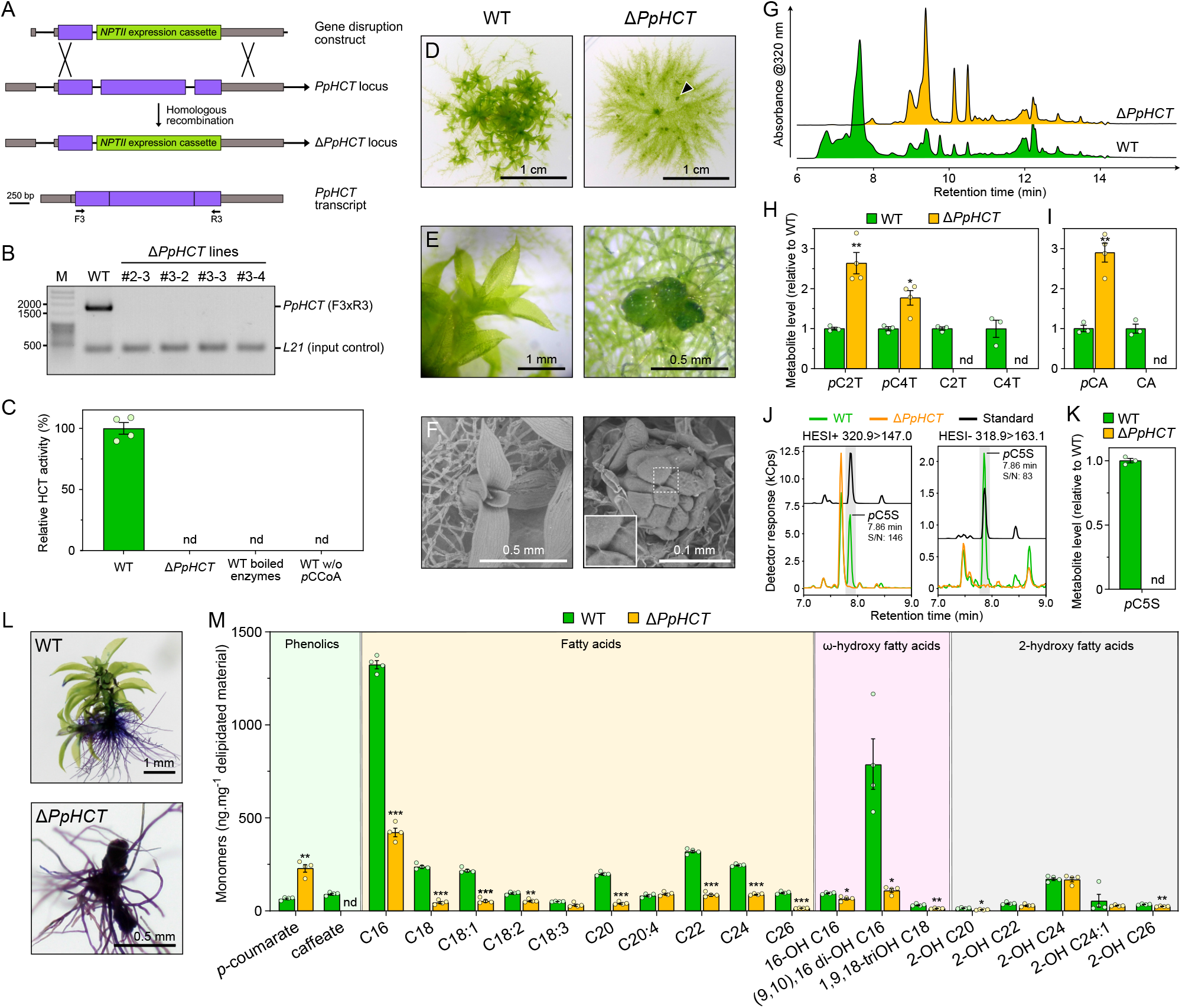
Investigation of *PpHCT* function *in planta*. Homologous recombination-mediated strategy for *PpHCT* gene disruption. A genomic fragment encompassing the second and third *PpHCT* exons was excised with simultaneous insertion of the *NPTII* selection cassette conferring resistance to G418. Binding sites of oligonucleotides used for characterization of the transgenic lines are shown. (B) Semi-quantitative RT-PCR analysis of the four Δ*PpHCT* KO lines confirms the absence of *PpHCT* transcripts. M, DNA size marker. (C) HCT activity in protein extracts from wild-type and Δ*PpHCT* gametophores. HCT activity was measured *in vitro* using shikimate and *p*-coumaroyl-CoA as substrates. Negative wild-type (WT) control assays involved boiled protein extracts or omission of *p*-coumaroyl-CoA (*p*CCoA). Results are the mean ± SEM of four independent enzyme assays, performed with protein extracts from each of the four independent Δ*PpHCT* mutant lines. nd, not detected. (D) Phenotype of four-week-old *P. patens* WT and Δ*PpHCT* colonies. Arrowhead points to a Δ*PpHCT* gametophore. (E) Magnified image of gametophores shown in (D). (F) SEM micrographs of four-week-old gametophores. For Δ*PpHCT*, inset shows intercellular adhesive structures. (G) Representative HPLC-UV chromatograms of WT and Δ*PpHCT* gametophore extracts. (H) Relative levels of phenolic threonate esters in gametophore extracts. *p*C2T, *p*-coumaroyl-2-*0*-threonate; *p*C4T, *p*-coumaroyl-4-*0*-threonate; C2T, caffeoyl-2-*0*-threonate; C4T, caffeoyl-4-*0*-threonate. (I) Relative levels of free hydroxycinnamic acids in gametophore extracts after acid hydrolysis. (J) Overlay of representative UHPLC-MS/MS chromatograms showing the absence of *p*-coumaroyl-5-*0*-shikimate (*p*C5S) in Δ*PpHCT* gametophore extracts. (K) Relative levels of *p*-coumaroyl-5-*0*-shikimate (*p*C5S) in gametophore extracts. Results are the mean ± SEM of three independent WT biological replicates and four independent Δ*PpHCT* mutant lines. (L) Toluidine blue staining of WT and a Δ*PpHCT* mutant. Protonema and rhizoids do not have a cuticle, and so are readily stained. (M) Compositional analysis of WT and Δ*PpHCT* gametophore cuticular biopolymers. Data are the mean ± SEM of four WT biological replicates and four independent Δ*PpHCT* mutant lines. WT vs mutant *t*-test adjusted P-value: *, P<0.05; **, P<0.01; ***, P<0.001.

### *PpHCT* deficiency impairs cuticle development

A previous analysis of a Δ*PpCYP98* mutant led us to conclude that the availability of caffeate, or a derivative, is required for normal *P. patens* gametophore development and cuticle formation (Renault et al, 2017). Since Δ*PpHCT* lines essentially phenocopied Δ*PpCYP98* at macroscopic and metabolic levels, we tested tissue surface permeability of mutant and WT gametophores using a toluidine blue staining assay, to assess for similar cuticle defects. The strong blue staining of the Δ*PpHCT* lines confirmed their increased surface permeability compared to WT (**Fig. 5L**, **Fig. S8**), consistent with reduced cuticle barrier properties associated with the *PpHCT* deletion. We also characterized the monomeric composition of the cuticular biopolymer from the Δ*PpHCT* gametophore and found differences in aliphatic or phenolic components compared with WT (**Fig. 5M**). The Δ*PpHCT* cuticle appeared to be devoid of caffeate residues, but showed a 3-fold increase in *p*-coumarate units compared with WT, consistent with the analysis of soluble phenolic compounds (**Fig. 5H, I**). This change in phenolic composition was accompanied by a substantial decrease in long-chain fatty acids (LCFA) and ω-hydroxylated LCFA, especially in the two most abundant monomers, palmitic acid (C16) and (9,10),16 di-hydroxypalmitic acid. A minor decrease in the total amounts of 2-OH-VLCFA (very-long chain fatty acids), derived from membrane sphingolipids (Molina et al., 2006), indicated that plasma membranes were only slightly affected, in contrast to the cuticular biopolymer (**Fig. 5M**).

### Conservation of HCT properties between bryophytes and angiosperms

Functional analysis of PpHCT suggested a conservation of *HCT* function over the ~500 million years that span embryophyte evolution. To provide support to this hypothesis, we first investigated *in vitro* the acyl acceptor permissiveness of recombinant HCT from *M. polymorpha* (MpHCT), which belongs to another major bryophyte group, and *A. thaliana* (AtHCT) (**Fig. 6A**). In contrast to PpHCT and MpHCT, AtHCT activity using threonate as an acyl acceptor was barely detectable (**Fig. 6A**). However, all three proteins had in common a preference for shikimate or quinate as an acceptor, indicative of conservation of HCT enzyme properties in embryophytes (**Fig. 6A**). To further assess the functional conservation of *HCT* genes across embryophyte evolution, we conducted transcomplementation experiments. The first step was to generate an *A. thaliana hct* null mutant since only RNA-interference lines, with residual *HCT* expression, were available. Following a CRISPR/Cas9-mediated strategy, we isolated a mutant allele characterized by a deletion of seven nucleotides in the *AtHCT* first exon, hereafter termed *hct^D^*^7^ mutant (**Fig. 6B**), which introduces a frameshift leading to a premature stop codon.

**Figure 6.**
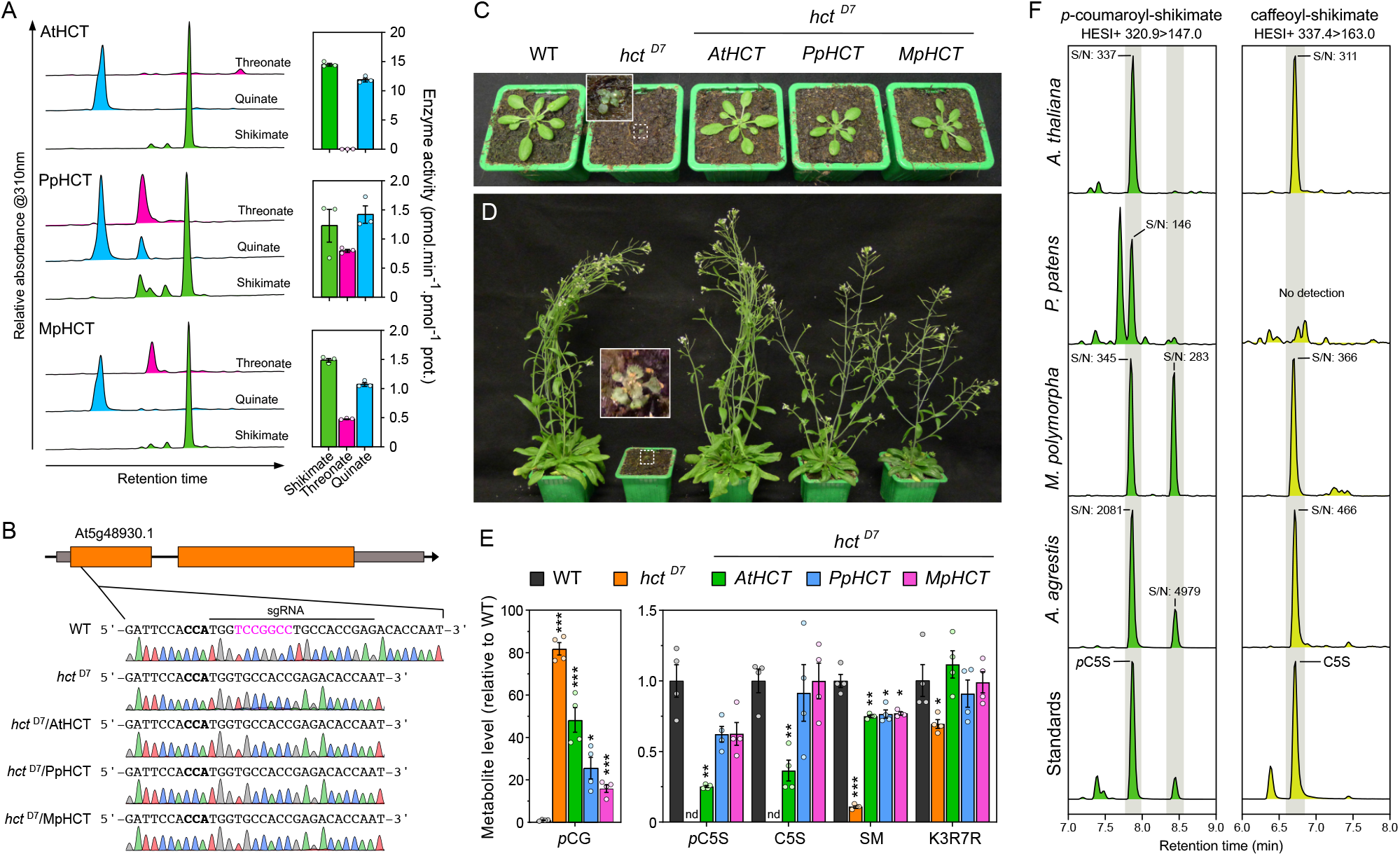
Evolutionary conservation of *HCT* function in embryophytes. (A) AtHCT, PpHCT and MpHCT acyl acceptor permissiveness was investigated using threonate, quinate or shikimate and *p*-coumaroyl-CoA as an acyl donor in *in vitro* end-point assays. Representative HPLC-UV chromatograms (left) and corresponding HCT activity (right) are shown for the different acyl acceptors. Results are the mean ± SEM of three independent enzyme assays. Note that the results for PpHCT are the same as those reported in **Fig. 3C**. (B) Schematic representation of the *AtHCT* locus and sequence of the CRISPR/Cas9 target site. The protospacer adjacent motif (NGG) is highlighted in bold. Sanger sequencing chromatograms of wild type (WT) and the homozygous *hct07* mutant transformed, or not, with *AtHCT*, *PpHCT* or *MpHCT* show the seven-nucleotide deletion in the *HCT* gene of *hct*D7 plants. (C-D) Phenotypes of three-week-old (C) and 60-day-old (D) *A. thaliana* wild type and the *hct07* null mutant transformed, or not, with *AtHCT*, *PpHCT* or *MpHCT* genes. (E) Relative levels of the phenolic esters *p*-coumaroyl-glucose (*p*CG), *p*-coumaroyl-5-*0*-shikimate (*p*C5S), caffeoyl-5-*0*-shikimate (C5S) and sinapoyl-malate (SM), and of the favonol kaempferol 3-*0*-rhamnoside 7-*0*-rhamnoside (K3R7R) in three-week-old rosettes. Results are the means ± SEM of four independent biological replicates. WT vs. mutant *t*-test adjusted P-value: *, P<0.05; **, P<0.01; ***, P<0.001. (F) Representative UHPLC-MS/MS chromatograms reporting the search for *p*-coumaroyl-5-*0*-shikimate and caffeoyl-5-*0*-shikimate in the bryophytes *P. patens*, *M. polymorpha* and *A. agrestis*, and the angiosperm *A. thaliana*. Additional positional isomers of *p*-coumaroyl-shikimate may occur in *M. polymorpha* and *A. agrestis*. S/N, signal-to-noise ratio.

We then transformed heterozygous *hct^D7^*^+/−^ plants with *AtHCT*, *PpHCT* and *MpHCT* coding sequences under control of the *A. thaliana C4H* promoter, which efficiently drives gene expression in phenylpropanoid-accumulating tissues (Weng et al., 2008, 2011; Sibout et al., 2016; Alber et al., 2019), and selected plants homozygous for both *hct^D7^* allele and complementation constructs (**Fig. 6B**). The *AtHCT* null mutation led to reduced growth (**Fig. 6C-D**), similar to previous observations of *HCT*-RNAi lines (Besseau et al., 2007; Li et al., 2010a), but this abnormal phenotype was entirely abolished by introducing an *HCT* coding sequence from *A. thaliana*, and almost completely in the case of *PpHCT* or *MpHCT* (**Fig. 6C-D**). Similar to *P. patens*, disruption of *HCT* function in Arabidopsis led to an increased permeability of aerial tissues to toluidine blue (**Tab. S10**). The *HCT* null mutation resulted in obvious changes in UV chromatograms (**Fig. S11**), which was confirmed by targeted analysis of diagnostic phenylpropanoid molecules. The targeted profiling revealed an 80-fold accumulation of *p*-coumaroyl-glucose in the *hct^D7^* mutant compared with WT, while *p*-coumaroyl and caffeoyl esters of shikimate were absent from the mutant (**Fig. 6E**). Residual levels of sinapoyl-malate, the main soluble phenolic ester in *A. thaliana* leaves, were detected in *hct^D7^* (~10% of wild-type level), possibly due to the alternative C3H/APX pathway using free *p*-coumarate, or promiscuous activities of C3’H on accumulating *p*-coumaroyl esters (e.g. *p*-coumaroyl-glucose; **Fig. 6F**). The level of the main *A. thaliana* leaf flavonoid, the flavonol kaempferol 3-*O*-rhamnoside 7-*O-*rhamnoside (Li et al., 2010a; Yin et al., 2014), was slightly reduced in *hct^D7^* compared with WT (**Fig. 6E**). This result does not match data from previous analyses of RNAi-*HCT* lines (Besseau et al., 2007; Li et al., 2010a), but is corroborated by HPLC-UV analysis indicating that no other phenylpropanoids, including flavonoids, over-accumulated in the *hct^D7^* null mutant to levels similar to wild-type sinapoyl-malate under our growth conditions (**Fig. S11**). All *hct^D7^* plant metabolic defects were, at least partially, complemented by transformation with *AtHCT*, *PpHCT* or *MpHCT* under the control of the *AtC4H* promoter (**Fig. 6E**). In particular, the ability to synthesize *p*-coumaroyl-5-*O*-shikimate and caffeoyl-5-*O*-shikimate was restored in all the *HCT*-complemented lines (**Fig. 6E**), consistent with functional conservation of bryophyte and angiosperm *HCT* genes. Threonate esters were not detected in any of the investigated Arabidopsis genotypes. To further establish phenolic shikimate esters as conserved metabolic intermediates during embryophyte evolution, we checked for their presence in representative species of the three major bryophyte lineages. In addition to *P. patens* and *A. thaliana* (**Fig. 6E-F, Fig. S12**), targeted analysis revealed the presence of *p*-coumaroyl-5-*O*-shikimate in the liverwort *M. polymorpha* and the hornwort *A. agrestis* (**Fig. 6F, Fig. S12**). With the exception of *P. patens*, the 3’-hydroxylated form of *p*-coumaroyl-5-*O*-shikimate, caffeoyl-5-*O*-shikimate, was detected in all plant samples. The results were consistently confirmed by both retention time comparison with molecular standards and simultaneous MS/MS analysis in positive and negative modes (**Fig. 6F, Fig. S12**). Parallel profiling of threonate esters in the same plant extracts suggested a lineage-specific pattern, since they were detected only in *P. patens* extracts (**Fig. S13**).

## DISCUSSION

The silencing of *HYDROXYCINNAMOYL-CoA:SHIKIMATE HYDROXYCINNAMOYL TRANSFERASE* in seed plants typically results in a strong reduction in the abundance, and/or compositional modification, of the biopolymer lignin, and is usually associated with stunted growth (Chen et al., 2006; Hoffmann et al., 2004; Wagner et al., 2007; Besseau et al., 2007; Gallego-Giraldo et al., 2011). Parallel *in vitro* and structural studies showed that tracheophyte HCTs consistently use shikimate as a preferred acyl acceptor to form *p*-coumaroyl-shikimate esters (Hoffmann et al., 2003; Chiang et al., 2018; Levsh et al., 2016; Lallemand et al., 2012), which in turn serve as substrates for C3’H enzymes (Schoch et al., 2001; Alber et al., 2019). Taken together, these data suggested a deep evolutionary conservation of HCT function in vascular plants. Here, through a multidisciplinary study of the bryophyte model *P. patens*, we are able to extend *HCT* functional conservation throughout the entirety of embryophyte evolution, pointing to an emergence in the last common ancestor of land plants, approximately 500 Ma (**Fig. 7A**). New methodologies have allowed us to refine our previous studies (Renault et al., 2017) and we conclude, based on new evidence, that shikimate esters are the native intermediates for phenolic ring 3-hydroxylation in *P. patens* (**Fig. 7B**). The absence of a *bona fide C3H/APX* gene and the presence of only distantly-related *CSE* homologs in *P. patens* (Ha et al., 2016; Barros et al., 2019) further supports a pivotal function of HCT in the moss phenylpropanoid pathway, for both producing and deconjugating shikimate esters (**Fig. 7B**).

**Figure 7.**
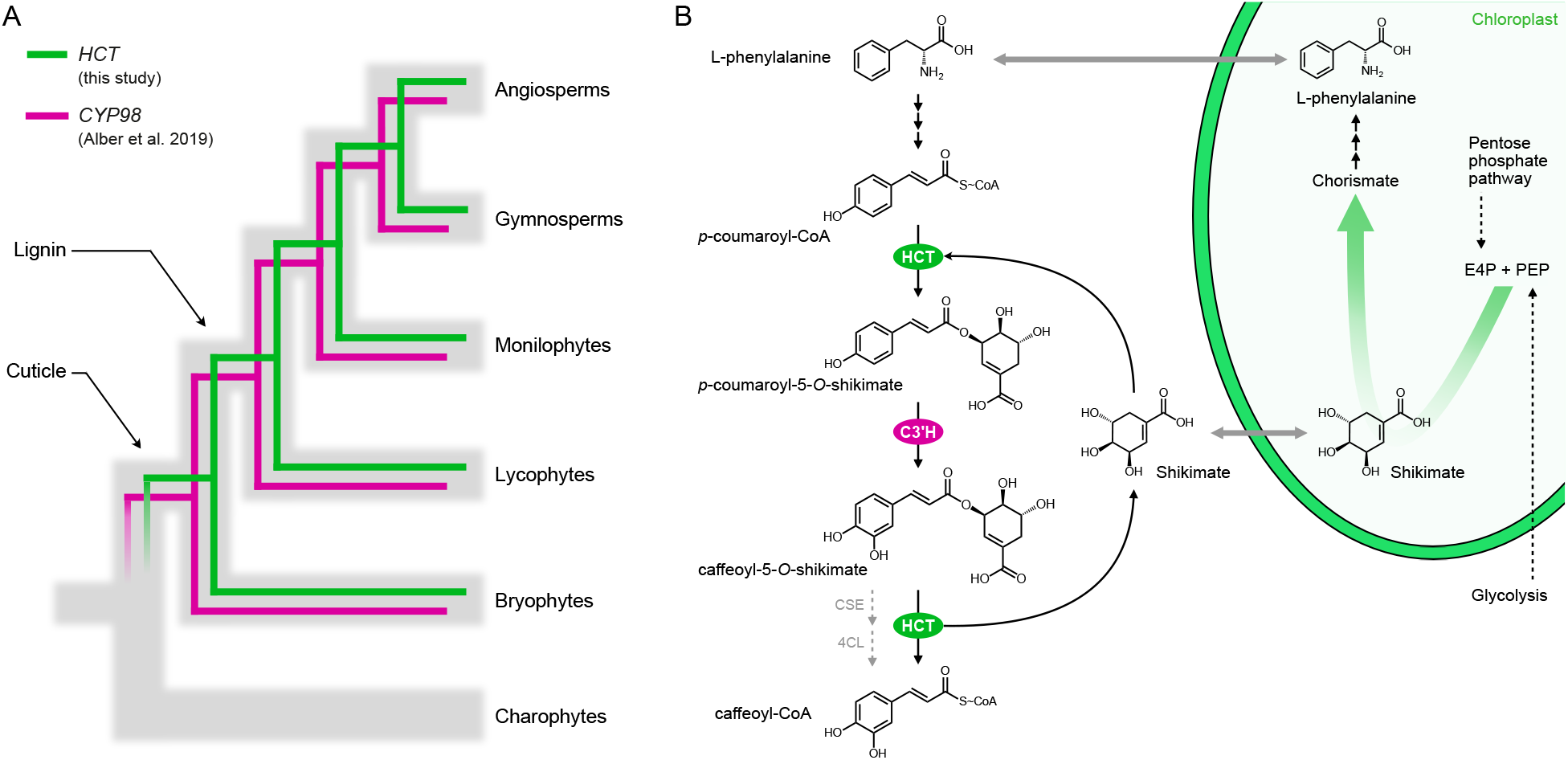
Evolutionary and metabolic models of HCT function. (A) Evolutionary model for the conservation of *HCT/CYP98* pair in embryophytes. Gray branches represent organismal evolution of species. Green and cyan branches represent *HCT* (this study) and *CYP98* (Alber et al., 2019) gene evolutions, respectively. *CYP98* encodes *p*-coumaroyl 3′hydroxylase (C3′H). (B) Metabolic model of *P. patens* phenylpropanoid pathway. HCT plays a pivotal role in controlling the formation and deconjugation of shikimate esters, the evolutionarily conserved intermediates for phenolic ring functionalization by C3′H enzyme. Shikimate partly derives from the pentose phosphate pathway and is a precursor of L-phenylalanine, potentially establishing a regulatory control of photosynthetic carbon allocation to the phenylpropanoid pathway through two critical steps. Occurrence of a CSE/4CL alternative route toward caffeoyl-CoA in *P. patens* remains an open question. E4P, D-erythrose 4-phosphate; PEP, phospho*eno/*pyruvate.

Our data also highlight a previously unappreciated complexity in the bryophyte phenylpropanoid pathway, which in *P. patens* produces both soluble esters and precursors of a hydrophobic apoplastic biopolymer. This metabolic typology is akin to that of flowering plants, which often produce lineage-specific soluble phenolic esters (e.g. sinapoyl-malate or chlorogenic acids), as well as essential precursors of biopolymers, such as monolignols. Soluble esters act as UV screens and antioxidants (Lehfeldt et al., 2000; Clé et al., 2008) and, as such, may be advantageous in particular ecological niches (Li et al., 2010b). We propose that threonate esters, which we found only in *P. patens*, are specialized stress-mitigating molecules, while the shikimate esters are evolutionarily conserved intermediates involved in phenolic ring functionalization. Threonate originates from the degradation of the plant-specific antioxidant ascorbate, possibly in the cell wall (Green and Fry, 2005), suggesting a connection between *P. patens* threonate ester biosynthesis and stress acclimation. Despite our efforts, we failed to identify the enzyme(s) responsible for threonate ester production in *P. patens*. Our preliminary data suggest that this neither involves a BAHD acyl transferase, nor a coenzyme A thioester acyl donor. In this respect, serine carboxypeptidase-like enzymes would be good candidates for catalyzing such a reaction using glucose esters as acyl donors, as is the case for sinapoyl-malate biosynthesis (Lehfeldt et al., 2000). On the other hand, evolutionary selection has led to the coupling of phenol-containing biopolymer biosynthesis with shikimate, a widespread molecule found in plants, bacteria and fungi. In plants, shikimate partly derives from the pentose phosphate pathway and is a precursor of aromatic amino acids, including L-phenylalanine (Maeda and Dudareva, 2012) (**Fig. 7B**). Cellular concentrations of shikimate are assumed to be lower than the *K*m of HCT enzymes for shikimate. It was therefore proposed that shikimate availability might serve as a biochemical mechanism to regulate photosynthetic carbon allocation to phenylpropanoid production, with HCT playing a key role in such a mechanism (Schoch et al., 2006; Adams et al., 2019) (**Fig. 7B**).

We here provide evidence that the *HCT* gene, and the associated shikimate-mediated regulation of the plant phenolic metabolism, appeared during plant terrestrialization in the last common ancestor of embryophytes, concomitant with the occurrence of a cuticle, but prior to lignin evolution (**Fig. 7A**). This evolutionary pattern matches that of *CYP98*, which encodes the downstream C3’H enzyme (Alber et al., 2019). A complex evolutionary interplay therefore likely shaped *HCT* and *CYP98* macro-evolutions and established the *HCT/CYP98* pairing as a core metabolic module within the phenylpropanoid pathway, deeply rooted in land plant evolution (**Fig. 7A**). The tight relationships between the two enzymes is further evidenced by their ability to physically interact and to form a supramolecular complex in *A. thaliana* (Bassard et al., 2012), a mechanism by which intermediate channeling could occur. Whether the phenylpropanoid pathway in *P. patens* is also organized at a supramolecular level however remains an open question. The *CYP98/HCT* pair also features lower scale evolutionary patterns, as illustrated by recurrent, independent duplications of the core *HCT* and *CYP98* genes, which led to the emergence of specialized phenolic compounds, such as rosmarinic acid and phenolamides (Matsuno et al., 2009; Liu et al., 2016; Levsh et al., 2019).

Both *PpCYP98* (Renault et al., 2017) and *PpHCT* knock-out mutants show stunted gametophore growth and organ fusion phenotypes, associated with a complete loss of cuticular caffeate units. Cuticles are essential to control water permeability, and provide plant protection against drought (Lü et al., 2012; Kosma et al., 2009) and other environmental stresses, including UV-B radiation (Krauss et al., 1997; Yeats and Rose, 2013). Thus, emergence of a cuticle with properties that enabled plant terrestrialization may have been dependent on the presence of a primordial phenylpropanoid pathway. The severe developmental defects of the Δ*PpHCT* and Δ*PpCYP98* mutants unfortunately prevent meaningful evaluation of their stress tolerance. Although found in substantial amounts in the cuticle of some tracheophytes, such as the leaf cuticle of *Solanum lycopersicum* (Bolger et al., 2014), hydroxycinnamic acids usually represent small proportions of the cuticle of vascular plants (Fich et al., 2016). The presence of large amounts of hydroxycinnamic acids might therefore be a typical, and possibly essential, feature of bryophyte lineages (Caldicott and Eglinton, 1976; Buda et al., 2013; Kong et al., 2020). Hydroxycinnamic acids might play an important role, since they are covalently attached to fatty acid monomers (Riley and Kolattukudy, 1975). We show here that the absence of caffeate in *P. patens* prevents the formation of the cuticle and cuticular biopolymer polymerization, as evidenced by the large decreases in the major cutin monomers C16 FA and (9,10),16 di-OH C16 FA in the Δ*PpHCT* lines, as was previously shown in the *PpCYP98* deletion mutants (Renault et al., 2017). A straightforward interpretation is that caffeate anchors, or shapes, the cuticle lipidic scaffold of *P. patens*. Such a function is apparently not fulfilled by *p*-coumarate, which accumulates in the Δ*PpHCT* cuticle. This might indicate an important role of phenolic ring functionalization for biopolymer formation, as is the case in natural plant lignins, which are predominantly derived from di- or tri-substituted phenolic precursors (Ralph et al., 2019). Whether the structural function of phenolic compounds in the cuticle is specific to bryophytes, or even *P. patens*, remains to be clarified. It was indeed shown that the absence of ferulate from *A. thaliana* cuticles did not noticeably reduce cuticle integrity (Rautengarten et al., 2012), while we report here an increased permeability to toluidine blue of the aerial tissues of Arabidopsis *hct^D7^* mutant. The enrichment of the *P. patens* cuticle with phenolic compounds potentially contributes various functional attributes, including UV protection, water/gas management, tissue scaffolding for erect growth and organ determination (i.e. organ fusion avoidance). We hypothesize that reduction of this bryophyte property was linked to the emergence of new, specialized biopolymers in tracheophytes, such as canonical lignin and suberin, which assumed some of the functions mediated by the phenol-enriched cuticle of bryophytes.

## METHODS

### Phylogenetic analysis

All BAHD sequences from *Physcomitrium patens* (moss, bryophyte), *Marchantia polymorpha* (liverwort, bryophyte), *Anthoceros agrestis* (hornwort, bryophyte), *Spirogloea muscicola* (*Zygnematophyceae*, charophyte), *Chara braunii* (*Charophyceae*, charophyte) and *Klebsormidium nitens* (*Klebsormidiophyceae*, charophyte) were retrieved by BLASTp search using AtHCT (At5g48930.1) as query (E-value<0.01). Truncated proteins with less than 420 residues were discarded. Obtained bryophyte and charophyte BAHDs were aligned with 34 functionally characterized BAHD protein (full list in **Tab. S1 and S2**) using the MUSCLE algorithm (Edgar, 2004) (alignment file available as **Dataset S1**). Ambiguous sites of the alignment were masked applying the Gblocks method (Castresana, 2000). Phylogenetic relationships were reconstructed with a maximum-likelihood approach using PhyML3.0 (Guindon et al., 2010). Selection of evolution model that best fits the dataset was guided by the SMS software; the tree was ultimately inferred from the LG +G+I+F model (Le and Gascuel, 2008). Initial tree(s) for the heuristic search were obtained automatically by applying the BioNJ algorithm, and by selecting the topology with superior log likelihood value. Best of nearest neighbor interchange (NNI) and subtree pruning and regrafting (SPR) methods were used for improving the tree. Branch tree supports were calculated with the approximate likelihood ratio test (Anisimova and Gascuel, 2006). Tree file is available as supplemental material (**Dataset S2**). Sequence manipulation was performed with Seaview 4 software (http://pbil.univ-lyon1.fr/) and phylogenetic analysis on PhyML server (http://www.atgc-montpellier.fr/phyml/).

### Homology modeling of proteins

3D models of *P. patens* (Pp3c2_29140), *C. braunii* (CHBRA170g00210) and *K. nitens* (kfl00513_0110) proteins were generated using the Modeler comparative module (Sali and Blundell, 1993) embedded in ChimeraX v1.0 software (Goddard et al., 2018) using *A. thaliana* HCT in complex with *p*-coumaroyl-5-*O*-shikimate (pdb entry: 5kju) as template. Prior to modeling, target proteins were aligned with embryophyte representative HCTs visible in **Fig. 1B** with MUSCLE algorithm (alignment file available as **Dataset S3**). Five models were automatically generated for each target proteins; 3D models with the best GA341 and zDOPE scores were kept for subsequent analyses. Potential hydrogen bonds linking protein residues and *p*-coumaroyl-5-*O*-shikimate were predicted with the ChimeraX *FindHBond* tool. Overlay and visualization of 3D protein models was performed with ChimeraX.

### Plant growth conditions

*Physcomitrella patens* (Hedw.) Bruch & Schimp., strain Gransden (IMSC acc. no. 40001, Lang et al., 2018) was cultured in liquid or on solid Knop medium (Reski and Abel, 1985) supplemented with 50 μmol.L^−1^ H3BO3, 50 μmol.L^−1^ MnSO4, 15 μmol.L^−1^ ZnSO4, 2.5 μmol.L^−1^ KI, 0.5 μmol.L^−1^ Na2MoO4, 0.05 μmol.L^−1^ CuSO4 and 0.05 μmol.L^−1^ CoCl2. Medium was solidified with 12 g.L^−1^ purified agar. *P. patens* gametophores were propagated on agar plates or in liquid cultures established by soft tissue disruption (~15 s). Liquid cultures were weekly subcultured and kept under constant agitation (130 rpm) for proper aeration. *Marchantia polymorpha* Tak-1 accession and *Anthoceros agrestis* Oxford accession were grown on half-strength Gamborg B5 medium solidified with 12 g.L^−1^ agar. Bryophytes were kept at 23°C under 16/8 h day/night cycle, light intensity set to 70 μmol.m^−2^.s^−1^. *Arabidopsis thaliana* plants (Col-0 background) were grown on soil, kept under 22/18°C, 16h/8h light/dark regime (100 μmol.m^−2^.s^−1^ light intensity) and were watered from the bottom every two days with tap water.

### Determination of gene expression by qRT-PCR

Total RNA was isolated from 10 mg of lyophilized tissue with 1 ml of TriReagent (Sigma-Aldrich). Samples were agitated 5 minutes at room temperature prior to centrifugation at 13,000 *g*, RT. After transfer of the supernatant to a new microtube, an equal volume of chloroform was added and samples were thoroughly vortexed and centrifuged at 13,000 *g* at RT to induce phase separation. The clear upper phase was recovered and transferred to a new microtube, total RNA was precipitated by adding 0.1 volume of sodium acetate (NaOAc, 3M, pH 5.2) and 2.5 volumes of absolute ethanol. After incubation at −20°C for 2h, RNA was spin down by centrifugation at 13,000 *g*, 4°C. Supernatant were discarded, the RNA pellet was washed with 1 ml of 70% ethanol, then dried at RT for 10 minutes. Total RNA was finally resuspended in DEPC-treated water. Twenty micrograms of RNA were treated with 5U of RQ1 DNaseI (Promega) and subsequently purified using phenol-chloroform (50/50, v/v) and precipitation by NaOAc/EtOH. One microgram of DNaseI-treated RNA was reverse-transcribed with oligo(dT) and the Superscript III enzyme (Thermo Scientific) in 20 μl reaction. Quantitative PCR reactions consisted of 10 ng cDNA, 500 nM of each primers and 5 μl of 2X LightCycler^®^ 480 SYBR Green I Master mix (Roche) in 10 μl final volume. Reactions were run in triplicates on a LightCycler^®^ 480 II device (Roche). The amplification program was 95 °C for 10 min and 40 cycles (95 °C denaturation for 10 s, annealing at 60 °C for 15 s, extension at 72 °C for 15 s), followed by a melting curve analysis from 55 to 95 °C to check for transcripts specificity. Crossing points (Cp) were determined using the manufacturer’s software. Cp values were corrected according to primer pair PCR efficiency computed with the LinReg PCR method (Ruijter et al., 2009). *Pp3c19_1800* and *Pp3c27_3270* genes were used as internal reference for expression normalization. List of qPCR primers is available in **Table S4**.

### GUS staining

Plant tissues were vacuum infiltrated during 10 min with X-Gluc solution (containing 50 mM potassium phosphate buffer pH 7.0, 0.5 mM ferrocyanide, 0.5 mM ferricyanide, 0.1% Triton X-100 1mM supplemented with 0.5 mg/mL X-Gluc) and incubated at 37 °C for 4.5 h. Chlorophyll was removed by washing tissues three times in 70% ethanol.

### Recombinant protein production

Cloning of AtHCT (*At5g48930*) coding sequence into the pGEX-KG vector and purification of the corresponding recombinant protein were performed as previously described (Hoffmann et al., 2003; Besseau et al., 2007). Coding sequences of PpHCT (*Pp3c2_29140*), MpHCT (*Mapoly0003s0277*) were PCR-amplified from *P. patens* Gransden and *M. polymorpha* Tak-1 cDNA respectively using Gateway-compatible primers (**Tab. S4**). The truncated PpHCT coding sequence, visible in **Fig. S4A,** was ordered as double-stranded gBlock (Integrated DNA Technologies) with Gateway compatible extensions. CDS were cloned into pDONR207 vector by BP Clonase reaction, then shuttled to the pHGWA expression vector by LR clonase reaction, allowing N-terminal fusion of protein with hexahistidine tag. *Escherichia coli* Rosetta2pLyS strain was transformed with recombined pHGWA plasmids and cultivated in ZYP-5052 autoinducible medium. Recombinant proteins were purified by immobilized metal affinity chromatography (IMAC) using an AKTA Pure 25 system equipped with HisTrap HP 1 mL column and submitted to gel filtration using a Superdex 200 increase 10/300 GL column (GE healthcare). Purified recombinant proteins were conserved at −80°C in 1x PBS solution containing 10% glycerol.

### *In vitro* enzyme assays

Five millimolar stock solutions of *p*-coumaroyl-CoA, caffeoyl-CoA and feruloyl-CoA (PlantMetaChem or MicroCombiChem) were prepared in H_2_O. Eighty millimolar stock solution of L-threonic acid was prepared from its hemicalcium salt (Sigma-Aldrich) in H2O containing 40 mM EDTA to chelate calcium and improve solubility. Fourty millimolar stock solutions of shikimate and D-quinate (Sigma-Aldrich) were prepared in H_2_O. Standard *in vitro* HCT assays were performed in 100 μL containing 50 mM potassium phosphate buffer pH 7.4, 1 mM dithiothreitol (DTT), 5 μg recombinant HCT protein, 5 mM acyl acceptor (-)-shikimate, D-quinate or L-threonate; Sigma-Aldrich) and 200 μM acyl-CoA. Reactions were initiated by addition of the acyl-CoA, incubated at 30°C for 25 minutes and stopped by addition of 100 μL acetonitrile. To determine PpHCT kinetic parameters, substrate and enzyme concentrations were optimized for each tested substrate. For shikimate, 50 ng protein, 200 μM *p*-coumaroyl-CoA and 0.125-8 mM (-)-shikimate were used. For quinate, 100 ng protein, 200 μM *p*-coumaroyl-CoA and 0.312-20 mM D-quinate were used. For threonate, 2 μg protein, 200 μM *p*-coumaroyl-CoA and 4-32 mM L-threonate were used. For *p*-coumaroyl-CoA, 50 ng protein, 8 mM shikimate and 12.5-400 μM *p*-coumaroyl-CoA, or 2 μg protein, 32 mM L-threonate and 12.5-600 μM *p*-coumaroyl-CoA were used. Reactions were initiated by addition of the saturating substrate, incubated at 30°C for 10 minutes and stopped by addition of 100 μL acetonitrile. Relative quantification of reaction products was performed by UHPLC-MS/MS. Absolute quantification of phenolic esters was performed on HPLC-UV with external calibration curves of corresponding free hydroxycinnamic acid (i.e. *p*-coumarate, caffeate and ferulate; Sigma-Aldrich). Kinetic parameters were calculated with nonlinear Michealis-Menten regression using the GraphPad Prism v4.8 software (**Fig. S5**). Threonate/shikimate competition assays were performed in 50 μl containing 50 mM potassium phosphate buffer pH 7.4, 1 mM DTT, 100 ng recombinant PpHCT, 2.5 mM L-threonic acid, 10 μM (-)-shikimic acid and 200 μM *p*-coumaroyl-CoA. Reactions were initiated by addition of the acyl-CoA, incubated for 15 min at 30°C and stopped with 50 μl methanol. Under these conditions, shikimate consumption was kept below 30%. Because of the complex nature of the competition assay, absolute quantification of reaction products *p*-coumaroyl-4-*O*-threonate and *p*-coumaroyl-5-*O*-shikimate was performed by UHPLC-MS/MS with external calibration curves of standard molecules. To test the PpHCT reverse reaction, *in vitro* assays were performed in 50 μl containing 50 mM potassium phosphate buffer pH 7.4, 1 mM DTT, 1 μg recombinant PpHCT, 250 μM coenzyme A (Sigma-Aldrich), 100 μM caffeoyl ester. Reactions were initiated by addition of the coenzyme A, incubated for 30 min at 30°C and stopped with 50 μl methanol. Absolute quantification of the reaction product caffeoyl-CoA was performed by HPLC-UV using external calibration curve of authentic molecule.

### Yeast metabolic engineering

For *P. patens* phenolic pathway reconstitution, *Pp4CL1, PpHCT, and PpCYP98* coding sequences were PCR-amplified from Gransden cDNA using Gateway-compatible primers (**Tab. S4**) and shuttled by LR recombination to yeast galactose-inducible expression vectors pAG424GAL, pAG423GAL and pAG425GAL (Alberti et al., 2007), respectively; *A. thaliana ATR1* coding sequence was PCR-amplified from Col-0 cDNA and transferred to pAG426GAL yeast expression vector. Recombined vectors were introduced in INVSc1 *S. cerevisiae* yeast strain (ThermoFisher Scientific) following the lithium acetate/polyethylene glycol method. Yeast transformant were selected on SC- media lacking relevant molecule(s) (6.7 g/L yeast nitrogen base without amino acids, 20 g/L glucose, appropriate concentration of relevant Yeast Synthetic Drop-out Medium; Sigma-Aldrich) and incubated three days at 30°C. For whole-cell metabolic assay, a 2.5 mL SC- liquid culture was inoculated with a yeast colony and incubated overnight at 180 rpm and 30°C. Cultures were centrifuged 5 min at 3,000*g* and cell pellets were washed in 25 mL sterile ultra-pure water and centrifuged again 5 min 3,000*g*. Cells were resuspended in 2.5 mL of liquid SC- medium supplemented with galactose instead of glucose to induce gene expression and incubated at 30°C, 180 rpm. Six hours after induction, yeast cultures were supplemented with 25 μL of 100 mM sterile *p*-coumarate solution in DMSO and 50 μL of 50 mM sterile L-threonic acid solution in water (5 mM final concentration each). Following substrates addition, cultures were incubated for 24 h at 30°C, 180 rpm. Metabolites were extracted from whole yeast cultures by adding one volume of methanol followed by thorough vortexing. Extracts were centrifuged at 16,000*g* for 10 min to spin down yeasts. Supernatants were recovered, dried *in vacuo* and resuspended in 50% methanol in 1/5 of initial volume. Concentrated extracts were analyzed by UHPLC-MS/MS.

### Generation of *P. patens* transgenic lines

Δ*PpHCT knock*-out mutants were generated by protoplast transfection with a genetic disruption construct allowing introduction of the *NPTII* expression cassette into *PpHCT* locus by homologous recombination. Genetic construct was made by assembling two 750 bp genomic regions PCR-amplified from *P. patens* genomic DNA with the *NPTII* selection cassette by GIBSON cloning. The assembled fragment was then PCR-amplified and blunt-end cloned into the pTA2 vector using the pJET1.2 cloning kit (ThermoFisher Scientific). *PpHCT* disruption construct was excised from vector backbone by *Eco*RI digestion, using restriction sites introduced by PCR. Final sterile DNA solution used for PEG-mediated protoplast transfection contained 45 μg of excised fragment in 0.1 M Ca(NO3)2. Protoplast isolation, transfection and regeneration were performed according to (Hohe et al., 2004). Transformants were selected on Knop plates supplemented with 25 mgl^−1^ geneticin (G418). For *PpHCT:uidA* reporter lines, two genomic regions for homologous recombination framing the *PpHCT* STOP codon were PCR-amplified from genomic DNA and assembled with the *uidA* reporter gene following the same procedures as described above. A linker sequence was introduced by PCR to limit GUS protein hindrance on PpHCT activity. The *PpHCT*:*uidA* construct was excised from vector backbone by *Nhe*I digestion. 50 μg of excised fragment were used for protoplasts transfection. Since *PpHCT:uidA* does not contain a selection marker, it was co-transfected with the pRT101 plasmid (Girke et al., 1998) containing the *NPTII* selection cassette. Transformants were selected on Knop plates supplemented with 25 mg l^−1^ geneticin (G418).

### Molecular characterization of *P. patens* transgenic lines

Proper genomic integration of DNA construct was assessed using a tailored PCR strategy (**Fig. S2, S7**) with primers listed in **Table S4**. Genomic DNA was extracted with DNA extraction buffer (75 mM Tris pH 8.8, 20 mM (NH4)2SO4 and 0.01 % Tween 20) during 15 min incubation at 45°C under agitation (1400 rpm). Two microliters were used for direct PCR using the Phire II DNA polymerase (Thermo Scientific) in a final volume of 20 μl. Δ*PpHCT* mutant lines with seamless 5’ and 3’ integration of the genetic construct at the desired locus were checked for the absence of full-length transcript. Total RNA was isolated and retrotranscribed as described above. *PpHCT* transcripts were amplified from two microliters of cDNA using the Phire II DNA polymerase (Thermo Scientific). The constitutively expressed *L21* gene (*Pp3c13_2360*), encoding a 60S ribosomal protein, was used as reference. Primers used for RT-PCR are listed in **Table S4**. Four transgenic lines with complete absence of *HCT* transcripts were selected for subsequent investigations. The MassRuler DNA Ladder Mix (ThermoFisher Scientific) was used as DNA size marker.

### Determination of HCT activity in *P. patens* protein extracts

Proteins were extracted from three-month-old WT and Δ*PpHCT* gametophores in 2 mL microtubes containing five volumes of extraction buffer (100 mM Tris-HCl pH 7.4, 10 % glycerol, 2 mM dithiothreitol, cOmplete™ EDTA-free Protease Inhibitor Cocktail). Samples were homogenized using 5 mm steel beads and a Tissuelyser II (Qiagen) operated at 30 Hz for 5 min. Following a centrifugation step (20,000*g*, 4°C, 40 min) supernatants were recovered and transferred to a 50 mL conical tube. Proteins were precipitated by slow addition to samples of ammonium sulfate up to 0.5 g/mL under constant agitation. Once ammonium sulfate was fully solubilized, samples were centrifuged for 20 min at 16,000*g* and 4°C, supernatants discarded and protein pellets resuspended in 5 mL of extraction buffer. A second round of precipitation and centrifugation was performed to fully remove plant endogenous metabolites. Protein pellets were resuspended in 500 μL of extraction buffer. Next, samples were centrifuged (5 min, 18,000*g*, 4°C) to pellet non-protein material, supernatants were transferred to new microtubes. Protein concentration was assessed with the Qubit Protein Assay Kit (ThermoFisher Scientific) and adjusted to 200 ng/μL with extraction buffer. All steps were performed at 4°C and samples were kept on ice. HCT activity in total proteins preparation was evaluated from 50 μl end-point enzyme assays containing 50 mM potassium phosphate buffer (pH 7.4), 2.5 μg total proteins, 1 mM dithiothreitol, 200 μM *p*-coumaroyl-CoA and 5 mM shikimate or threonate. Reactions were initiated by addition of *p*-coumaroyl-CoA, incubated at 30°C and stopped after 1 hour by addition of 50 μL acetonitrile. Production of *p*-coumaroyl-shikimate was monitored by UHPLC-MS/MS. Relative HCT activity was computed from *p*-coumaroyl-shikimate peak area and expressed as a percentage of WT.

### Plant tissue collection and metabolite extraction

Liquid cultured gametophores were harvested five weeks after the last tissue disruption, and one week after nutrient medium change. Plant material was collected by filtration on a 100 μm pore size sieve, quickly blotted on paper towel and snap-frozen in liquid nitrogen. For *M. polymorpha* and *A. agrestis*, one-month-old thalli were harvested from Petri plates and snapped-frozen in liquid nitrogen. For *A. thaliana*, whole 3-week-old rosettes were harvested from soil-grown plants and snap-frozen in liquid nitrogen. Samples were lyophilized for two days; dry material was homogenized using 5 mm steel beads and a Tissuelyser II (Qiagen) for 1 min at 30 Hz. Metabolites were extracted from 8 mg dry plant powder following a methanol:chloroform:water protocol as described previously (Renault et al., 2017), except that 500 μl methanol, 250 μl chloroform and 500 μl water were used. For shikimate ester detection in *P. patens*, 250 μl of metabolic extracts were dried in vacuo, dry residues were resuspended in 50 μl 50% methanol before analysis. Acid hydrolysis of metabolic extract was conducted as reported before (Renault et al., 2017).

### HPLC-UV chromatography

Metabolite separation and detection were carried out on a high-performance liquid chromatography system (Alliance 2695; Waters) coupled to a photodiode array detector (PDA 2996; Waters). Ten to twenty microliters were injected onto Kinetex Core-Shell C18 column (100 x 4.6 mm, 2.6 μm particle size or 150 x 4.6 mm, 5 μm particle size; Phenomenex). Needle and injection loops were successively washed with weak (95% water/5% acetonitrile) and strong (20% water/80% acetonitrile) solvents. For phenolics, the mobile phase consisted of a mix of HPLC grade water (A) and acetonitrile (B), both containing 0.1% formic acid. The elution program was as follows: 0.0 min - 95% A; 15.0 min, 5% A (curve 8); 17.0 min, 5% A (curve 6); 18.0 min, 95% A (curve 6); 20.0 min, 95% A. The flow was set to 1 ml/min and column temperature to 35°C. For acyl-CoA analysis, the mobile phase consisted of a mix of 20 mM sodium phosphate pH 5.3 prepared in HPLC grade water (A) and acetonitrile (B). The elution program was as follows: 0.0 min - 95% A; 1.0 min, 95% A; 9.0 min, 50% A; 10 min, 40% A; 10.5 min, 5% A; 12.0 min, 5% A. The flow was set to 1 ml/min and column temperature to 35°C and the absorbance was recorded between 250 and 400 nm. Data were processed with the Empower 3 Software (Waters).

### Targeted metabolic profiling by UHPLC-MS/MS

Separation and detection of metabolites were carried out on an ultra-high-performance liquid chromatography (UHPLC; Dionex UltiMate 3000, ThermoFisher Scientific) coupled to an EvoQ Elite LC-TQ (MS/MS) mass spectrometer equipped with a heated electrospray ionization source (HESI) (Bruker). Nitrogen was used as the drying (30 L/h flow rate) and nebulizing gas (35 L/h flow rate). The interface temperature was set to 350°C and the source temperature to 300°C. The capillary voltage was set to 3.5 kV both for positive and negative ionization modes. MS data acquisition and LC piloting were performed with the Bruker MS Workstation 8 and Compass Hystar 4.1 SR1 softwares, respectively. Metabolites were ionized in either positive and negative modes and detected using specific MRM methods (**Table S5**). Bruker MS Data Review software was used to integrate peaks and report corresponding areas. For phenolic molecules, three microliters of sample were injected onto a C18 Cortecs^©^ UPLC^©^ T3 column (150 × 2.1 mm, 1.6 μm; Waters) and eluted with a mix of LC-MS grade water (A) and acetonitrile (B), both containing 0.1% formic acid to keep molecules in protonated form. After each injection, the needle and injection loop were washed with 25% acetonitrile solution. The elution program was as follows: 0.0 min - 5% B; 1.0 min - 5% B; 11.5 min - 100% B (curve 8); 13.0 min - 100% B; 14.0 min - 5% B (curve 6); 15.0 min - 5% B; total run time: 15 min. Flow was set to 0.400 mL/min and column temperature to 35°C. Metabolite peak area was normalized to plant dry weight; metabolite level was expressed relative to WT. For central polar metabolites, two microliters of same plant extracts used for phenolics analysis were injected onto an Acquity UPLC^©^ HSS PFP column (150 × 2.1 mm, 1.8 μm; Waters) and eluted with a mix of LC-MS grade water (A) and acetonitrile (B), both containing 0.1% formic acid. After each injection, needle and injection loop were washed with 25% acetonitrile solution. Elution program was as follows: 0.0 min - 0% B; 2.0 min - 0% B; 5.0 min - 25% B; 10.0 min - 35% B; 10.5 min - 95% B; 12.5 min - 95% B; 12.6 min - 0% B; 15.0 min - 0% B; total run time: 15 min. Flow was set to 0.250 mL/min and column temperature to 40°C. The absolute level of polar central metabolites was calculated according to plant dry weight and external calibration curves of authentic molecules and expressed as μmoles of compounds per g of plant dry weight (μmol.g^−1^ DW).

### Cuticular biopolymer compositional analysis

Cutin monomers analysis was performed on the same gametophore samples used for metabolic analysis, following a previously published protocol (Renault et al., 2017). Briefly, tissues were delipidated by extensive washing with a series of solvents. The delipidated tissues, including cuticle, were dried, weighed and chemically treated (12:3:5 methanol: methyl acetate: 25% sodium methoxide, 60°C, o/n) to depolymerize cutin. Released monomers were then derivatized with pyridine and BSTFA (N,Obis(trimethylsilyl)trifluoroacetamide), dried again by heating under a stream of nitrogen, and resuspended in 100 μl of chloroform. The samples were analyzed by gas chromatography (GC) using an Agilent GC 6850 with a Flame Ionization Detector. Compounds were identified based on a comparison of retention times with standards, and by performing GC–mass spectrometry (MS) using an Agilent GC 6890 coupled to a JEOL GC MATE II mass spectrometer. Monomer levels were normalized to internal standards and dry delipidated tissue weights.

### Permeability assay

The permeability test was performed by immersing gametophores in a 0.05% toluidine blue solution for two minutes, then rinsing with water until the washing solution was clear.

### Production and complementation of an Arabidopsis *hct* null mutant

We generated an *hct* null mutant by CRISPR/Cas9-mediated gene inactivation as described earlier (DiGennaro et al., 2018). Briefly, *BbsI* restriction enzyme was used to introduce into the At-psgR/GW plasmid a double strand fragment resulting from 5’-GATTGCTCGGTGGCAGGCCGGACCA and 5’-AAACTGGTCCGGCCTGCCACCGAGC oligonucleotides annealing, which targets *HCT* region CTCGGTGGCAGGCCGGACCATGG. At-psgR/GW with *HCT* genomic target were transferred into the pEarleyGate 301 vector by LR Clonase reaction (ThermoFisher Scientific). The recombined pEarleyGate 301 vector was introduced into *Agrobacterium tumefaciens* GV3101 and used to transform Arabidopsis Col-0 by the floral dip method. Genotyping of T1 and T2 plants was performed by PCR amplifying the genomic sequence spanning the *HCT* target site using 5’-CCTTCTGAGAGAGTTGGTCGAC and 5’-CTAGCTCGGAGGAGTGTTCG oligonucleotides, followed by *AvaII* restriction digestion and run on an agarose gel to assess restriction site loss. The loss of the *At-psgR/GW* cassette at T2 or subsequent generation was assessed by sensitivity to selective agent (glyphosate). A line, free of the *AtpsgR/GW* cassette and harboring a 7 bp deletion 28 bp after the initiation codon was isolated for this study and named *hct^D7^*. Mutation at the desired locus was confirmed by Sanger sequencing using PCR fragment generated with 5’-CCTTCTGAGAGAGTTGGTCGAC and 5’-CTAGCTCGGAGGAGTGTTCG oligonucleotides. *hct^D7^* was subsequently used for transcomplementation assays with *AtHCT*, *PpHCT* and *MpHCT* coding sequences. To this end, Gateway pENTRY vectors harboring coding sequences were recombined with the binary pCC0996 vector that contains a 2977 bp promoter fragment from *A. thaliana C4H* gene (Weng et al., 2011). Resulting plant expression vectors were introduced into *Agrobacterium tumefaciens* GV3101 and used to transform heterozygous *hct^D7^* plants by the floral dip method. Transformants were selected with BASTA and *hct^D7^* allele was monitored along the selection process as described above. Experiments were performed with T3 plants homozygous for both the mutant allele and the transcomplemention construct.

### Replication and statistical analyses

Unless otherwise noted, independent biological replicates correspond to pooled plants from an independent container. For *P. patens*, one replicate derived from plants grown in one flask; for *M. polymorpha* and *A. agrestis*, one replicate derived from plants grown in one Petri plate; for *A. thaliana*, one replicate derived from plants grown in one pot. All statistical analyses were performed with GraphPad v8 software. For enzyme catalytic parameters, 95% confidence intervals were computed from nonlinear regression curves based on three independent enzyme assays. For metabolic profiling data, multiple two-tailed unpaired Student *t*-tests were performed to compare wild-type and mutant means; *P*-values were corrected using the Holm-Šídák method. Results from statistical analyses are shown in **Tables S6-9**.

## SUPPLEMENTAL DATA

**Supplemental Figure S1.** Multiple sequence alignment of protein region containing residues critical for HCT activity.

**Supplemental Figure S2.** Molecular characterization of *PpHCT:uidA* reporter lines.

**Supplemental Figure S3.** Multiple sequence alignment of representative embryophyte HCTs.

**Supplemental Figure S4.** Sequence, multiple sequence alignment and catalytic properties of a trunctated PpHCT protein.

**Supplemental Figure S5.** Saturation curves of PpHCT activity used to infer kinetic parameters.

**Supplemental Figure S6.** *In vitro* threonate/shikimate competition assay.

**Supplemental Figure S7.** Assessment of the catalytic function *K. nitens* HCT homologous protein kfl00513_0110 in yeast.

**Supplemental Figure S8.** Molecular and phenotypic characterization of Δ*PpHCT* mutant lines.

**Supplemental Figure S9.** Search for *P. patens* enzymes able to produce *p*-coumaroyl-threonate from *p*-coumaroyl-CoA.

**Supplemental Figure S10.** Toluidine blue staining assay of Arabidopsis *hct^D7^* mutant.

**Supplemental Figure S11.** UV fingerprinting of metabolic extracts from *hct^D7^* lines.

**Supplemental Figure S12.** Shikimate ester occurrence in bryophytes as evidenced by HESI UHPLC-MS/MS analysis in negative mode.

**Supplemental Figure S13.** Threonate esters occur in *P. patens* only as evidenced by HESI+ UHPLC-MS/MS analysis.

**Supplemental Table S1.** List of functionally characterized hydroxycinnamoyl-CoA-dependent BAHD acyltransferases used for the phylogenetic analysis.

**Supplemental Table S2.** List of uncharacterized BAHD acyltransferases used for the phylogenetic analysis.

**Supplemental Table S3.** Absolute level of L-phenylalanine, L-malate, D-quinate, L-threonate and (-)-shikimate in plant tissues.

**Supplemental Table S4.** List of oligonucleotides used in the study.

**Supplemental Table S5.** List of multiple reaction monitoring methods used for metabolite analysis.

**Supplemental Table S6.** Results of t-test from Figure 5H.

**Supplemental Table S7.**Results of t-test from Figure 5I.

**Supplemental Table S8.**Results of t-test from Figure 5M.

**Supplemental Table S9.**Results of t-test from Figure 6E.

**Supplemental Dataset S1.** Multiple protein alignment of BAHD protein sequences used for phylogeny reconstruction.

**Supplemental Dataset S2.** BAHD phylogenetic tree file.

**Supplemental Dataset S3.** Multiple protein alignment used to reconstruct HCT tridimensional models.

## ACKNOWLEDGEMENTS

This work received support from the initiative of excellence IDEX-Unistra (ANR-10-IDEX-0002-02, HR), the Agence Nationale de la Recherche (ANR-19-CE20-0017, HR), the grant-in-aid “Diversity of Biological Mechanisms” from the Institut des Sciences Biologiques – CNRS (DBM2020, HR), the Deutsche Forschungsgemeinschaft (DFG, German Research Foundation) under Germany’s Excellence Strategy (EXC-2189 – Project ID: 390939984, RR), the National Science Foundation (NSF-1517546, JKCR) and the Agriculture and Food Research Initiative of the United States Department of Agriculture (2016-67013-24732, J.K.C.R.). Authors would like to thank Pr. Takayuki Kohchi (Kyoto University) and Dr. Isabel Monte (University of Zürich) for providing *Marchantia polymorpha* and *Anthoceros agrestis* plants, respectively. We are grateful to Annette Alber who performed initial *in vitro* PpHCT enzyme assays.

## AUTHORS CONTRIBUTIONS

LK and HR designed the research; LK, SK, EG, KT, DG, IS, LH, JZ and HR performed research; LK and HR analyzed data; LK and HR wrote the manuscript with critical input of JKCR, RR and DW.

