## Supplemental Figures and Tables for "Function of the HYDROXYCINNAMOYL-CoA:SHIKIMATE HYDROXYCINNAMOYL TRANSFERASE is evolutionarily conserved in embryophytes"

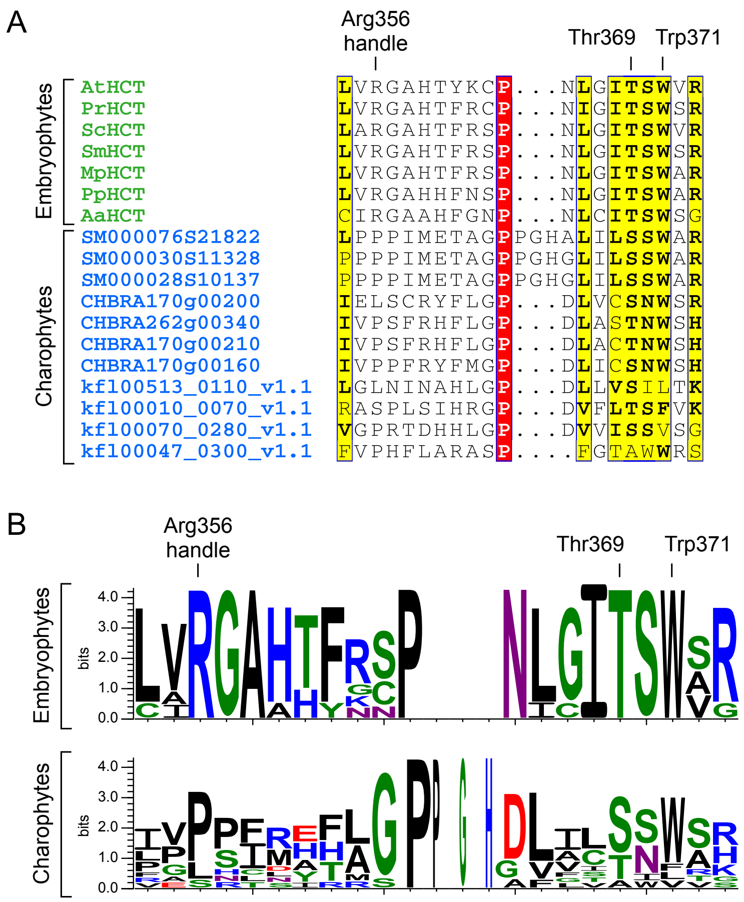


#### **Supplemental Figure S1**. **Multiple sequence alignment of protein region containing residues critical for HCT activity.**

(A) Sequences from representative embryophyte HCTs were aligned with all charophytes BAHDs. Positions that are identical are highlighted with a red background; positions with >70% similarity are highlighted with a yellow background. Alignment was performed with MUSCLE (Edgar, 2004) and prepared using the ESPript 3.0 server (Robert and Gouet, 2014). Residues critical for HCT catalytic activity are indicated and numbered according to AtHCT. (B) Graphical representation of sequence conservation in bryophytes and charophytes using Weblogo 3 server (Crooks et al., 2004). Supports Figure 1.

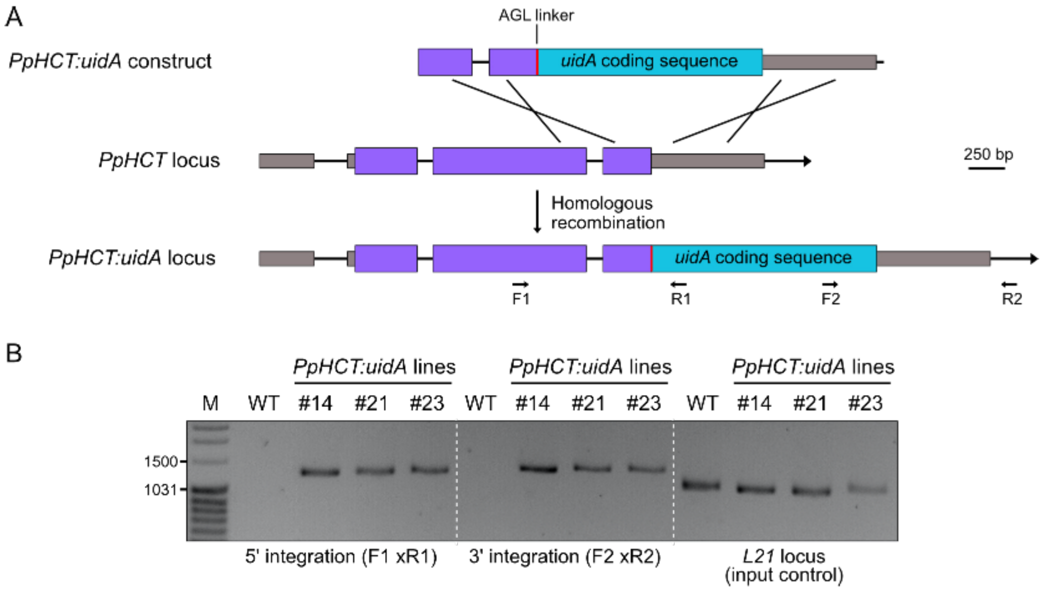


**Supplemental Figure S2. Molecular characterization of *PpHCT:uidA* reporter lines.**

(A) Homologous recombination-mediated strategy for *PpHCT* fusion with the *uidA* reporter gene. The *uidA* gene preceded by an alanine-glycine-leucine (AGL) linker sequence was inserted, substituting *PpHCT* STOP codon. (B) Agarose gel picture reporting PCR validation of correct integration of the construct in *PpHCT* genomic locus of the three G418-selected transgenic lines. Oligonucleotide hybridization sites are indicated in (A). M, DNA size marker (MassRuler DNA Ladder Mix, ThermoFisher Scientific). Supports Figure 2.

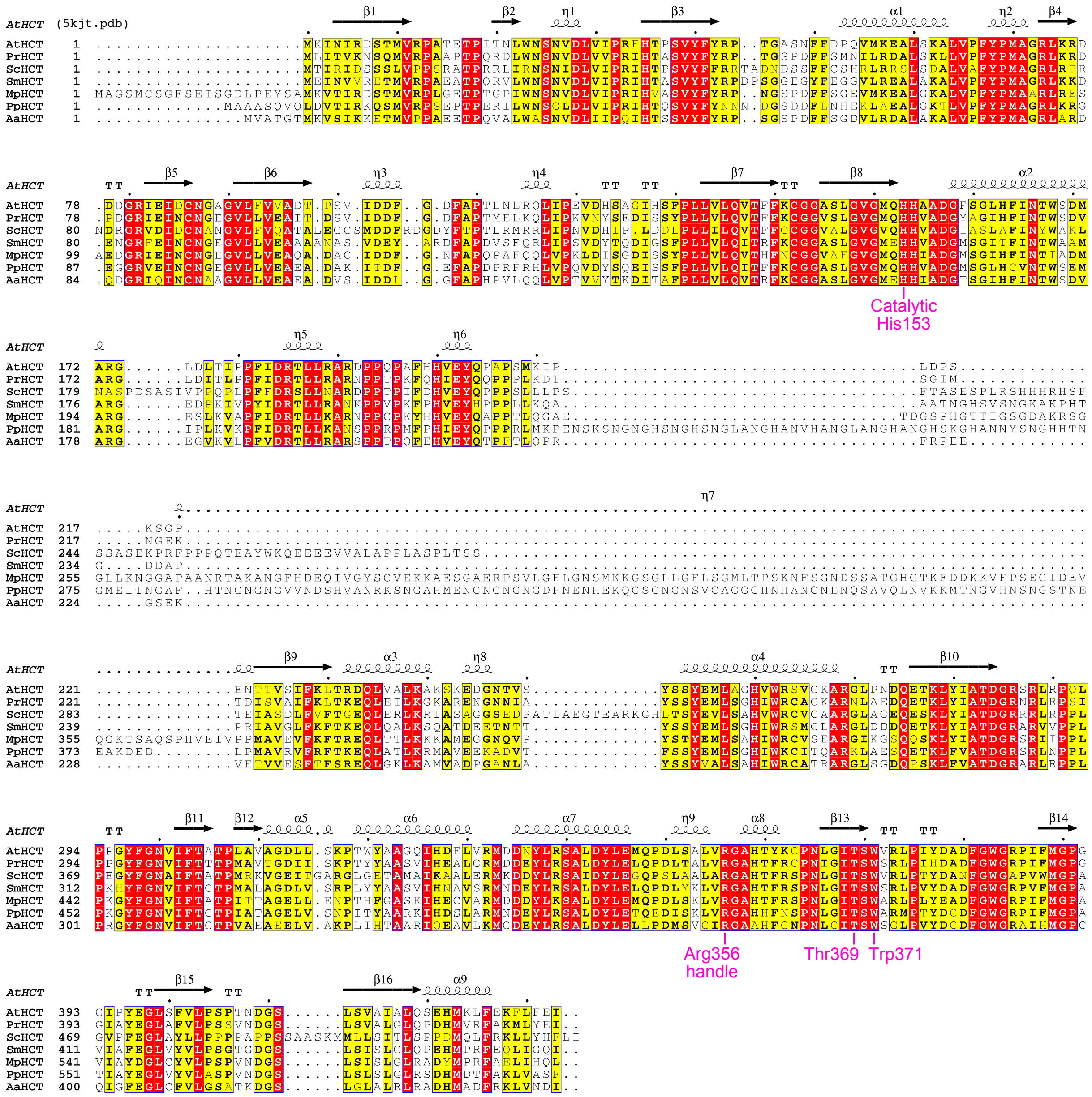


**Supplemental Figure S3**. **Multiple sequence alignment of representative embryophyte HCTs.**

Positions that are identical are highlighted with a red background; positions with >70% similarity are highlighted with a yellow background. Alignment was performed with MUSCLE (Edgar, 2004) and prepared using the ESPript 3.0 server (Robert and Gouet, 2014). Protein data bank entry for AtHCT is 5kjt. Residues important for HCT catalytic activity are indicated and numbered according to AtHCT. Supports Figures 1 and 3.

A

```
>PpHCT_truncated
ATGGCCGCCGCAAGTCAAGTTCAGTTGGATGTGACCATTAGGAAGCAGTCCATGGTGCGTCCCAGTGAACCGACACCAGAGAGAATACTCTGG
AACTCAGGTTTGGACCTTGTCATTCCTAGAATCCACACGCAGAGCGTGTATTTTTACAACAACAATGATGGAAGTGACGATTTCTTGAACCAT
GAGAAGCTTGCGGAGGCGCTGGGCAAGACATTGGTGCCATTTTACCCCATGGCTGGCCGACTGAAACGGGGGGAAGGAGGAAGAGTCGAGATT
AACTGTAATGGAGAAGGAGTGCTTCTTGTCGAAGCTGAAGCAGATGCTAAGATTACTGATTTCGGCGAGTTTGCTCCAGACCCCCGTTTTCGC
CACCTCGTTCCTCAAGTGGACTACTCCCAAGAGATATCTTCGTTTCCGCTTCTGGTTCTCCAGATCACGTTCTTCAAATGCGGTGGTGCGTCG
CTAGGAGTGGGCATGCAGCATCACGTTGCAGATGGCATGTCTGGCCTTCACTGCGTCAACACATGGTCGGAGATGGCGAGAGGGATCCCTCTC
AAAGTTAAACCGTTTATTGACCGCACGTTGTTGAAGGCGAACTCCCCTCCTAGACCCATGTTTCCTCACATTGAGTACCAACCTCCGCCCAGA
TTAATGAAACCTGAGAATTCTGCTAAAGACGAGGACCTGCCCATGGCTGTGCGTGTCTTCCGATTCACCAAAGAGCAATTGGCCACTTTGAAA
CGAATGGCAGTGGAGGAGAAGGCCGATGTGACCTTCAGCAGCTACGAAATGCTGTCAGGCCACATCTGGAAATGCATCACGCAAGCTCGGAAG
TTGGCAGAGTCTCAAGAAACCAAGCTTTTCGTCGCAACCGACGGCCGGTCGCGGCTCAATCCGCCACTTCCGAAGGGCTACTTCGGCAATGTG
ATCTTCACTTGCACTCCTATTGCAACGGCTGGGGAGTTGGTATCAAATCCAATAACATATGCTGCGAGAAAGATTCATGATTCGTTGGCGAGA
ATGAATGATGAATATTTGCGATCTGCACTTGACTACCTGGAGACGCAAGAGGATATCAGCAAACTCGTCCGCGGCGCACACCACTTCAATTCT
CCAAACCTGGGGATCACCTCCTGGGCGCGAATGCCCACTTACGACTGCGATTTTGGCTGGGGGAGACCGATTTTCATGGACCAGCTACAATA
GCCTATGAAGGATTGGTGTATGTGTTAGCCAGCCCTGTCAACGACGGAAGCTTGAGCCTGTCCTTGGGCTTGAGATCCGACCACATGGACACA
TTTGCCAAACTAGTGGCATCCTTCTAA
```

B

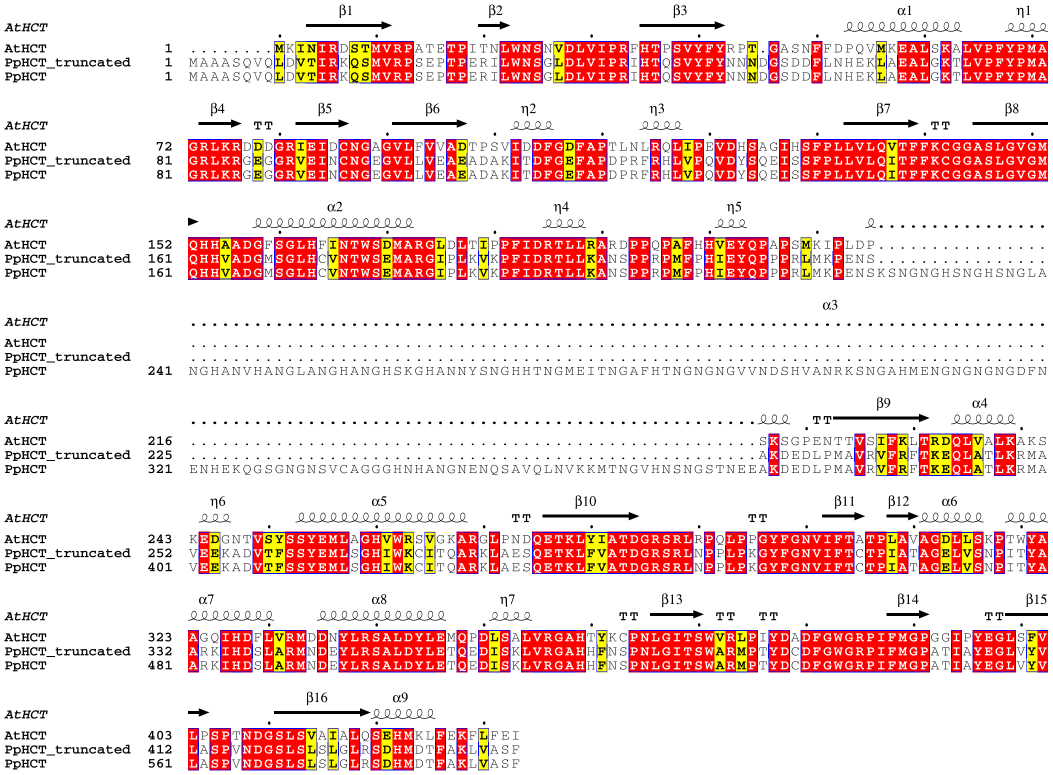


C

| **Acyl-acceptor** | **Protein** | **Activity** (pmol.min$^{-1}$.pmol$^{-1}$ prot.) |
|---|---|---|
| Threonate | PpHCT | 0.80 ±0.03 |
| | PpHCT_truncated | 0.49 ±0.01 |
| Quinate | PpHCT | 1.43 ±0.15 |
| | PpHCT_truncated | 1.49 ±0.03 |
| Shikimate | PpHCT | 1.24 ±0.28 |
| | PpHCT_truncated | 1.15 ±0.03 |

**Supplemental Figure S4. Sequence, multiple sequence alignment and catalytic properties of a trunctated PpHCT protein.**

(A) Sequence of PpHCT truncated (PpHCT_truncated) from its central loop (position 225 to 373). (B) Multiple sequence alignment of AtHCT with PpHCT and its truncated version. Positions that are identical are highlighted with a red background; positions with >70% similarity are highlighted with a yellow background. Alignment was performed with MUSCLE (Edgar, 2004) and prepared using the ESPript 3.0 server (Robert and Gouet, 2014). Protein data bank entry for AtHCT is 5kjt. (C) PpHCT and PpHCT_truncated activities with the combination of *p*-coumaroyl-CoA and one of the three acyl acceptors quinate, threonate and shikimate. Enzyme activity was calculated based on end-point assays analyzed by HPLC-UV. Results are the means ± SEM of three independent enzyme assays. Note that results for PpHCT are the same to those reported in Fig. 3C. Supports Figure 3.

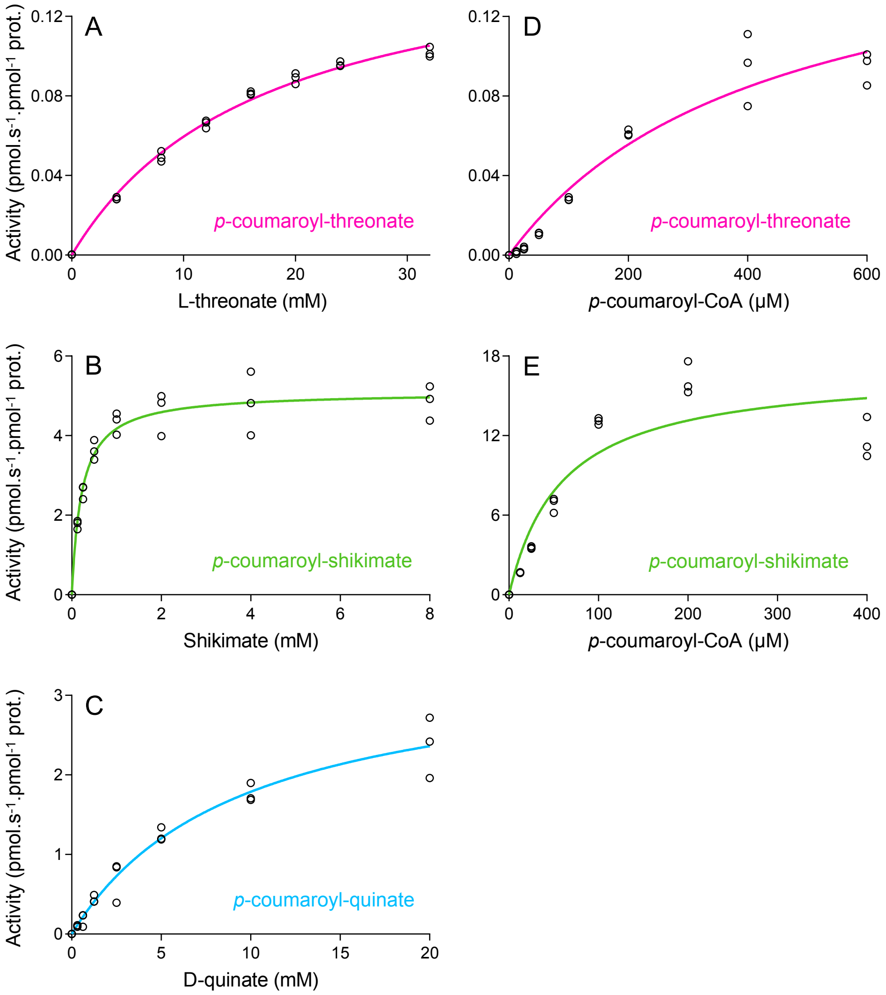


**Supplemental Figure S5. Saturation curves of PpHCT activity used to infer kinetic parameters.**

(A-C) Saturation curves of PpHCT activity, expressed as pmoles of product per second per pmoles of enzyme, using *p*-coumaroyl-CoA as fixed substrate and threonate (A), shikimate (B) or quinate (C) as varying substrate. (D-E) PpHCT activity saturation curves using *p*-coumaroyl-CoA as varying substrate and threonate (D) or shikimate (E) as fixed substrate. Michaelis–Menten nonlinear regression curves are visible. Results are means of three independent enzyme assays. Supports Table 1.

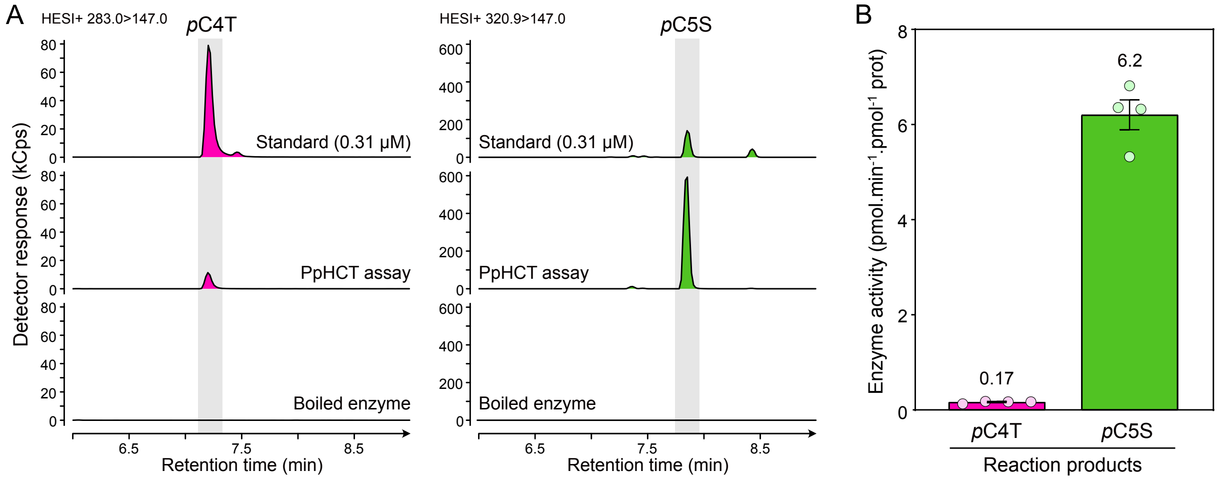


### Supplemental Figure S6. *In vitro* threonate/shikimate competition assay.

Recombinant PpHCT was incubated with 200 μM *p*-coumaroyl-CoA, 2.5 mM L-threonate and 10 μM (-)-shikimate. Concurrent control assays were performed with boiled enzyme. (A) UHPLC-MS/MS chromatograms showing the simultaneous production of *p*-coumaroyl-4-*O*-threonate (*p*C4T) and *p*-coumaroyl-5-*O*-shikimate (*p*C5S) by PpHCT *in vitro*. (B) PpHCT activity leading to *p*C4T (threonate acylation) or *p*C5S (shikimate acylation) production. Enzyme activity was calculated based on end-point assays analyzed by UHPLC-MS/MS. Results are means ± SEM of four independent enzyme assays. Supports Table 1.

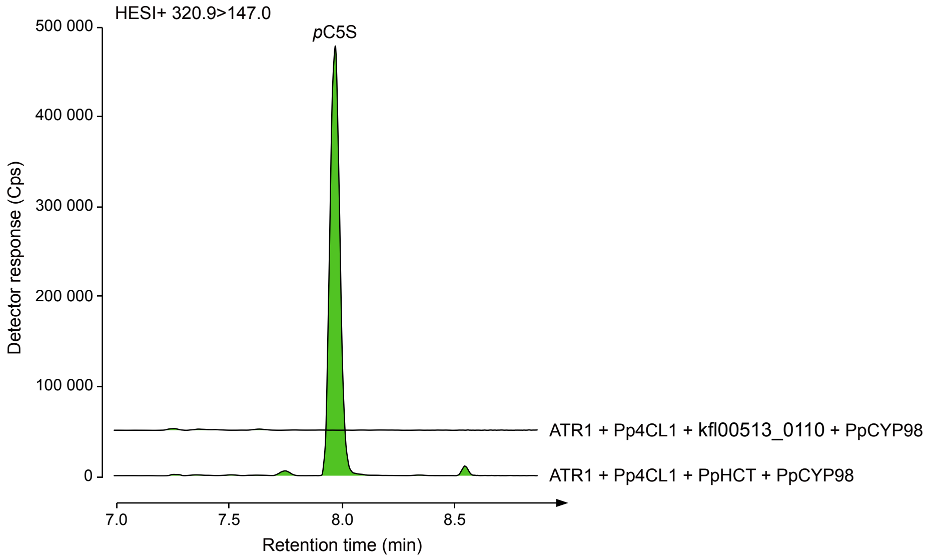


### Supplemental Figure S7. Assessment of the catalytic function *K. nitens* HCT homologous protein kfl00513_0110 in yeast.

*K. nitens HCT* homologous gene *kfl00513_0110* was expressed in *S. cerevisiae* along with *Pp4CL1*, *PpCYP98* and *ATR1*. After protein production step, yeast cultures were supplemented with *p*-coumarate and L-threonate. Targeted analysis *p*-coumaroyl-5-*O*-shikimate in yeast extracts was performed by HESI+ UHPLC-MS/MS. Yeast whole-cell assay with *PpHCT* was used as positive control. Chromatogram Y axes are linked. *p*C5S, *p*-coumaroyl-5-*O*-shikimate. Supports Figures 1 and 3.

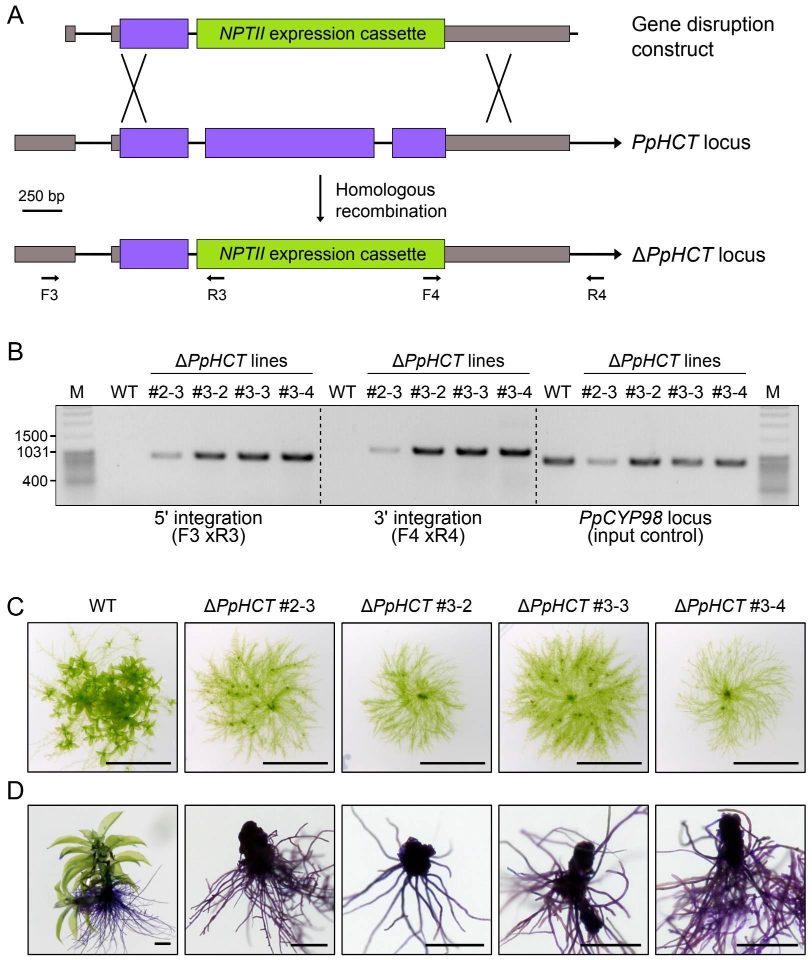


**Supplemental Figure S8. Molecular and phenotypic characterization of Δ*PpHCT* mutant lines.**

(A) Homologous recombination-mediated strategy for *PpHCT* gene disruption. A genomic fragment encompassing the second and third *PpHCT* exons was excised and replaced by the *NPTII* selection cassette conferring resistance to G418. (B) Agarose gel picture reporting PCR validation of correct integration of disruption construct in *PpHCT* genomic locus of the four G418-selected mutant lines. Oligonucleotide hybridization sites are indicated in (A). M, DNA size marker (MassRuler DNA Ladder Mix, ThermoFisher Scientific). (C) Phenotype of four-week-old *P. patens* WT and Δ*PpHCT* mutant colonies. Scale bars, 1 cm. (D) Toluidine blue staining assay of four-week-old WT and Δ*PpHCT* plants. Scale bars, 0.5 mm. Supports Figure 5.

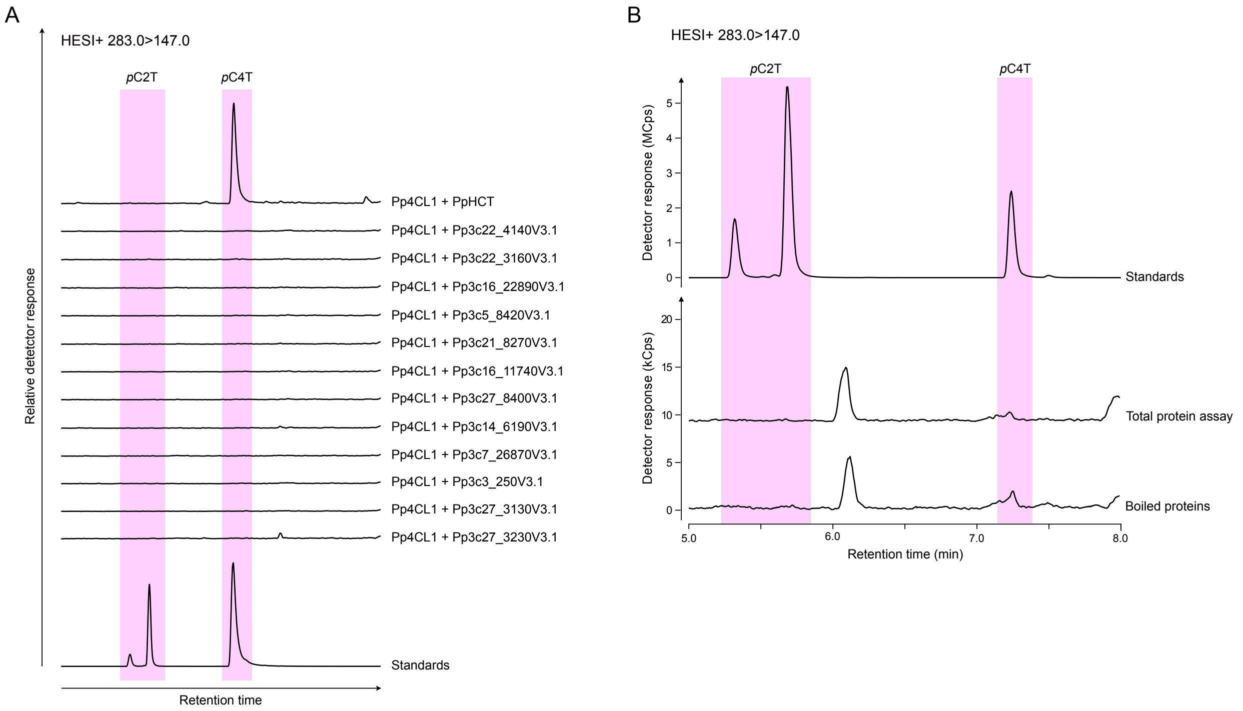


**Supplemental Figure S9. Search for *P. patens* enzymes able to produce *p*-coumaroyl-threonate using *p*-coumaroyl-CoA as an acyl donor.**

(A) All BAHD acyltransferases from *P. patens* identified via the Pfam motif PF02458 were cloned from gametophore cDNA and shuttled to a galactose-inducible yeast expression vector by Gateway cloning. *BAHDs* were individually expressed in the yeast *S. cerevisiae* together with *Pp4CL1*. Following galactose-induced protein production, yeast cultures were supplemented with *p*-coumarate and L- threonate and allow to grow further. *p*-coumaroyl-threonate esters were analyzed in yeast culture extracts by HESI+ UHPLC-MS/MS. Whole-cell yeast assay with *PpHCT* was used as positive control. Note that Pp3c27_3140 and Pp3c24_11730, which encode BAHD acyltransferases, were not included in the screen since they could not be PCR-amplified from gametophyte cDNA, in line with publicly available expression data (Perroud et al., 2018). Y axes of BAHD-related chromatograms are linked. (B) Total protein extract from wild-type *P. patens* gametophore was assayed for *p*-coumaroyl-threonate production *in vitro* (see Methods section for details). Concurrent negative control was performed with boiled total protein. *p*C2T, *p*-coumaroyl-2-*O*-threonate; *p*C4T, *p*-coumaroyl-4-*O*-threonate. Supports Figure 5.

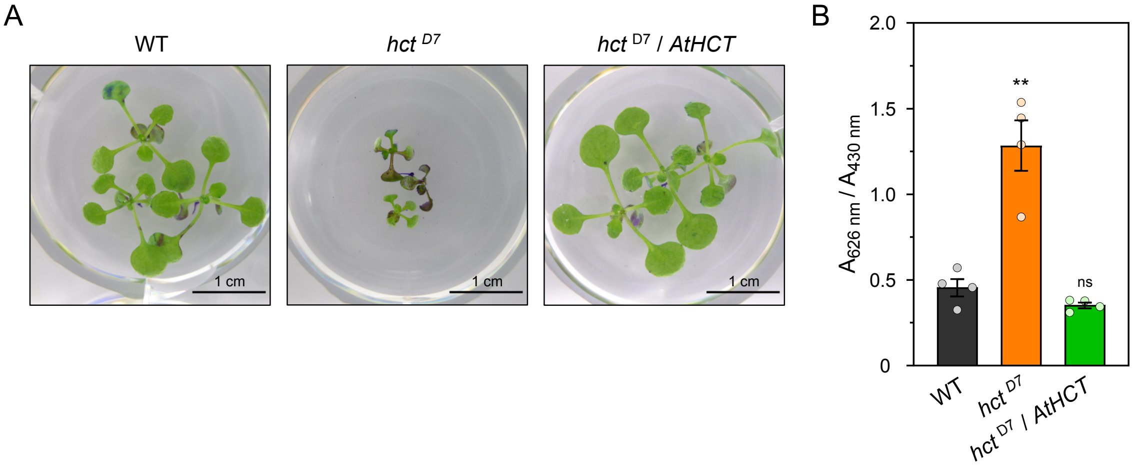


**Supplemental Figure S10. Toluidine blue staining assay of Arabidopsis *hct*$^{D7}$ mutant.**

Three-week-old plants were immersed for 5 min in 0.05% toluidine blue solution, rinsed with water until solution was clear, imaged and finally freeze-dried for quantitative analysis of staining. (A) Pictures of wild-type, *hct*$^{D7}$ mutant and *hct*$^{D7}$ complemented with *AtHCT* coding sequence after toluidine blue staining assay. (B) Quantitative analysis of toluidine blue staining. After pigments extraction according to Xu et al. (2020), absorbance at 626 nm and 430 nm was recorded and served for calculation of the $A_{626\ nm}/A_{430\ nm}$ ratio that accounts for staining intensity. Data are the mean ± SEM of four biological replicates consisting each of one plant (WT and *hct*$^{D7}$/*AtHCT*) or three plants (*hct*$^{D7}$). WT vs. mutant *t*-test adjusted P-value: **, P<0.01; ns, not significant. Supports Figure 6.

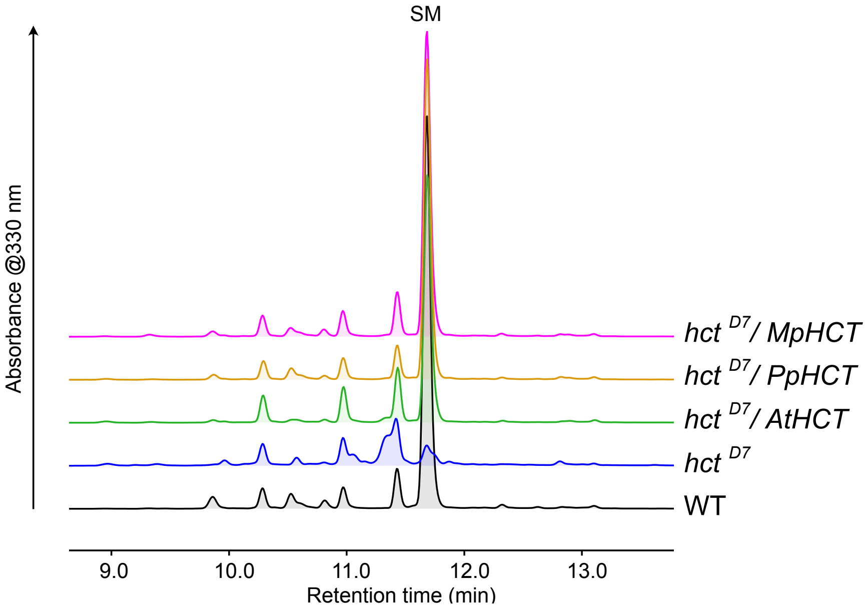


**Supplemental Figure S11. UV fingerprinting of metabolic extracts from *hct*$^{D7}$ lines**.

Metabolite extracts from three-week-old *A. thaliana* rosettes were analyzed by HPLC-UV. Shown are representative chromatograms of WT and *hct*$^{D7}$ mutant complemented, or not, with *AtHCT*, *PpHCT* and *MpHCT* coding sequences. SM, sinapoyl-malate. Supports Figure 6.

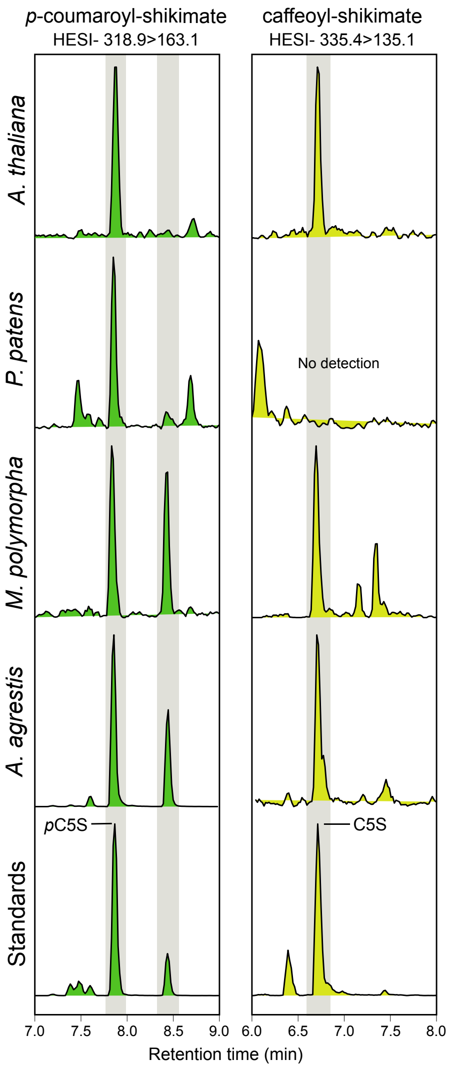


**Supplemental Figure S12. Shikimate ester occurrence in bryophytes as evidenced by HESI UHPLC-MS/MS analysis in negative mode.**

Occurrence of *p*-coumaroyl-5-*O*-shikimate (*p*C5S) and caffeoyl-5-*O*-shikimate (C5S) in the bryophytes *P. patens*, *M. polymorpha* and *A. agrestis*, and the angiosperm *A. thaliana* was confirmed using HESI- MRM methods. Note that caffeoyl-5-*O*-shikimate could not be detected in *P. patens* gametophore extracts. Results were observed in at least three independent samples. Supports Figure 6.

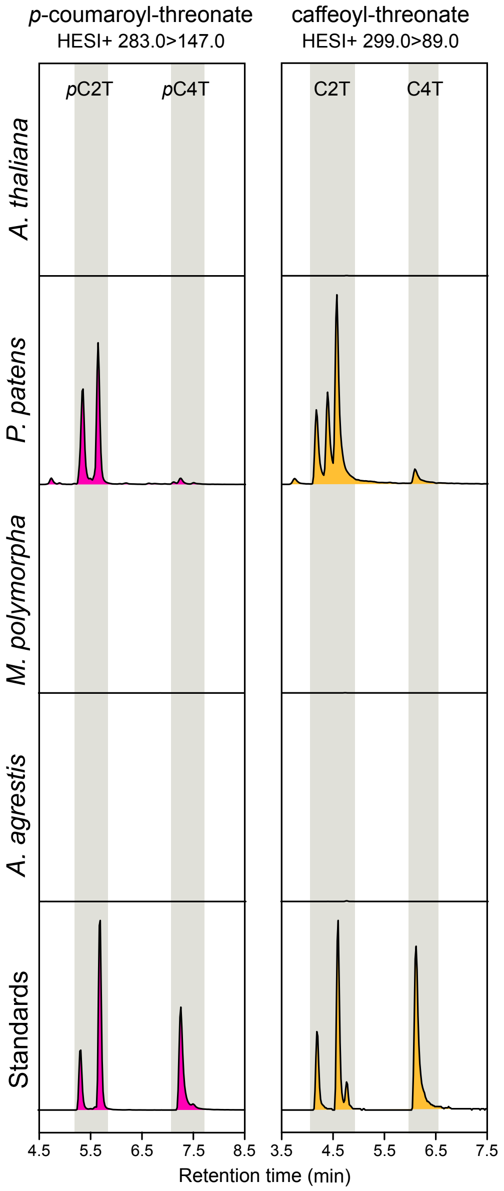


**Supplemental Figure S13. Threonate esters occur in *P. patens* only as evidenced by HESI+ UHPLC-MS/MS analysis.**

Occurrence of *p*-coumaroyl-2-*O*-threonate (*p*C2T), *p*-coumaroyl-4-*O*-threonate (*p*C4T), caffeoyl-2-*O*-threonate (C2T) and caffeoyl-4-*O*-threonate (C4T) in the bryophytes *P. patens*, *M. polymorpha* and *A. agrestis*, and the angiosperm *A. thaliana* was checked by HESI+ MRM methods. For plant chromatograms, Y axes are linked. Results were observed in at least three independent samples. Supports Figure 6.

### SUPPLEMENTAL TABLES

**Supplemental Table S1. List of functionally characterized hydroxycinnamoyl-CoA-dependent BAHD acyltransferases used for the phylogenetic analysis.**

| Preferred acceptor | Protein name | Species | Accession number |
|---|---|---|---|
| Shikimate | AtHCT | *Arabidopsis thaliana* | AT5G48930 |
| | CbHCT | *Coleus blumei* | CBI83579.1 |
| | CcHCT | *Coffea canephora* | ABO47805.1 |
| | CsHCT | *Cucumis sativa* | NP_001295843.1 |
| | MtHCT | *Medicago truncatula* | XP_013454529.1 |
| | NtHCT | *Nicotiana tabacum* | XP_009615699.1 |
| | PrHCT | *Pinus radiata* | ABO52899.1 |
| | PtHCT1 | *Populus trichocarpa* | ACC63882.1 |
| | PtHCT6 | *Populus trichocarpa* | XP_006368492.1 |
| | PvHCT1a | *Panicum virgatum* | AFY17066.1 |
| | TpHCT1 | *Trifolium pratense* | ACI16630.1 |
| Shikimate/Glycerol | OsHCT4 | *Oryza sativa* | NP_001057003 |
| Quinate | CcaHQT1 | *Cynara cardunculus* | ACF37072.1 |
| | CcaHQT2 | *Cynara cardunculus* | ACJ23164 |
| | CcHQT | *Coffea canephora* | ABO77956.1 |
| | EcHQT1 | *Erythroxylum coca* | AFF19203.1 |
| | LjHQT | *Lonicera japonica* | ACZ52698 |
| | NtHQT | *Nicotiana tabacum* | CAE46932.1 |
| | SlHQT | *Solanum lycopersicum* | NP_001234850.1 |
| | StHQT | *Solanum tuberosum* | ABA46756.1 |
| Aliphatics | AtASFT | *Arabidopsis thaliana* | NP_851111.1 |
| | AtDCF | *Arabidopsis thaliana* | AAQ62868.1 |
| | AtFACT | *Arabidopsis thaliana* | NP_201161.1 |
| | PtFHT1 | *Populus trichocarpa* | XP_002298644.2 |
| | StFHT | *Solanum tuberosum* | NP_001275190.1 |
| Amines | AtSHT | *Arabidopsis thaliana* | NP_179497.1 |
| | HCBT | *Dianthus caryophyllus* | O23917.1 |
| | HvACT | *Hordeum vulgare* | AAO73071.1 |
| | NaAT1 | *Nicotiana attenuata* | AET80688.1 |
| 4-hydroxyphenyllactate | CbRAS | *Coleus blumei* | A0PDV5.1 |
| | LaAAT1 | *Lavandula angustifolia* | ABI48360.1 |
| | MoRAS | *Melissa officinalis* | G0LD36.1 |
| Aromatic alcohols | PtSABT | *Populus trichocarpa* | XP_002319767 |
| Malate | TpHCT2 | *Trifolium pratense* | ACI16631.1 |

**Supplemental Table S2. List of uncharacterized BAHD acyltransferases used for the phylogenetic analysis.**

| Species | Gene Id |
|---|---|
| *Anthoceros agrestis* (bryophyte, hornwort) | AagrBONN_117.3413.1 |
| | AagrBONN_117.3458.1 |
| | AagrBONN_228.4525.1 |
| | AagrBONN_344.1108.1 |
| | AagrBONN_344.2322.1 |
| | AagrBONN_344.3712.1 |
| | AagrBONN_344.4211.2 |
| | AagrBONN_344.4213.1 |
| | AagrBONN_344.4620.1 |
| | AagrBONN_344.5246.1 |
| | AagrBONN_344.5370.1 |
| | AagrBONN_362.2723.1 |
| | AagrBONN_362.324.1 |
| | AagrBONN_362.596.1 |
| | AagrBONN_368.3175.2 |
| *Marchantia polymorpha* (bryophyte, liverwort) | Mapoly0001s0569.1 |
| | Mapoly0002s0087.1 |
| | Mapoly0003s0198.1 |
| | Mapoly0003s0277.1 |
| | Mapoly0005s0261.1 |
| | Mapoly0007s0149.1 |
| | Mapoly0008s0162.1 |
| | Mapoly0010s0137.1 |
| | Mapoly0010s0138.1 |
| | Mapoly0011s0054.1 |
| | Mapoly0012s0077.1 |
| | Mapoly0015s0010.1 |
| | Mapoly0015s0042.1 |
| | Mapoly0027s0021.1 |
| | Mapoly0027s0022.1 |
| | Mapoly0045s0065.1 |
| | Mapoly0045s0128.1 |
| | Mapoly0048s0047.1 |
| | Mapoly0088s0075.1 |
| | Mapoly0095s0065.1 |
| | Mapoly0107s0010.1 |
| | Mapoly0107s0011.1 |
| | Mapoly0107s0013.1 |

| | |
|---|---|
| | Mapoly0118s0027.1 |
| | Mapoly0124s0023.1 |
| | Mapoly0133s0035.1 |
| | Mapoly0157s0023.1 |
| | Mapoly0161s0008.1 |
| | Mapoly0306s0001.1 |
| *Physcomitrium patens* (bryophyte, moss) | Pp3c14_6190V3.1 |
| | Pp3c16_11740V3.1 |
| | Pp3c16_22890V3.1 |
| | Pp3c2_29140V3.1 |
| | Pp3c21_8270V3.1 |
| | Pp3c22_3160V3.1 |
| | Pp3c22_4140V3.1 |
| | Pp3c24_11730V3.1 |
| | Pp3c27_3130V3.1 |
| | Pp3c27_3230V3.1 |
| | Pp3c27_8400V3.1 |
| | Pp3c3_250V3.1 |
| | Pp3c5_8420V3.1 |
| | Pp3c7_26870V3.1 |
| *Spirogloea muscicola* (charophyte, *Zygnematophyceae*) | SM000028S10137 |
| | SM000030S11328 |
| | SM000076S21822 |
| *Chara braunii* (charophyte, *Characeae*) | CHBRA170g00160 |
| | CHBRA170g00200 |
| | CHBRA170g00210 |
| | CHBRA262g00340 |
| *Klebsormidium nitens* (charophyte, *Klebsormidiophyceae*) | kfl00010_0070_v1.1 |
| | kfl00047_0300_v1.1 |
| | kfl00070_0280_v1.1 |
| | kfl00513_0110_v1.1 |

**Supplemental Table S3. Absolute level of L-phenylalanine, L-malate, D-quinate, L-threonate and (-)-shikimate in plant tissues.**

Metabolites were extracted from three-week-old *Arabidopsis thaliana* rosettes, and one-month-old *Anthoceros agrestis*, *Marchantia polymorpha* and *Physcomitirum patens* gametophytes and analyzed by UHPLC-MS/MS. Absolute levels were calculated using external calibration curves of authentic molecules and expressed as µmoles of molecule per gram of plant dry weight (µmol.g$^{-1}$ DW). nd, not detected.

| Metabolite level (µmol.g$^{-1}$ DW) | *A. thaliana* | *A. agrestis* | *M. polymorpha* | *P. patens* |
|---|---|---|---|---|
| Phenylalanine | 0.28 ±0.06 | 0.88 ±0.29 | 0.23 ±0.01 | 0.89 ±0.11 |
| Malate | 20.9 ±2.3 | 14.9 ±0.3 | 25.5 ±1.1 | 16.7 ±0.7 |
| Quinate | nd | nd | nd | nd |
| Threonate | 0.11 ±0.01 | 0.01 ±0.06 | 0.04 ±0.06 | 2.22 ±0.06 |
| Shikimate | 0.061 ±0.006 | 0.025 ±0.001 | 0.43 ±0.03 | 0.009 ±0.001 |
| Threonate/Shikimate | 1.8 ±0.19 | 0.5 ±0.05 | 0.1 ±0.001 | 257 ±7 |

### Supplemental Table S4. List of oligonucleotides used in the study.

Nucleotides in lower case were involved in the cloning process, nucleotides in red indicate restriction sites, nucleotides in blue were neither specific to target sequence nor involved in cloning.

| Log id | Name | Sequence (5' > 3') |
|---|---|---|
| *PpHCT:uidA* construct (GIBSON cloning) | | |
| LK0316 | HR1_HCT_GUS_fwd | actcactatagggcgctagcGGACCTGCCCATGGCTGT |
| LK0317 | HR1_HCT_GUS_rev | ttaagcctgcGAAGGATGCCACTAGTTTGGCAAATG |
| LK0318 | GUS_gibson_F_HCT | ggcatccttcGCAGGCTTAATGTTACGTC |
| LK0319 | GUS_gibson_R_HCT | gatgaagactTCATTGTTTGCCTCCCTG |
| LK0320 | HR2_HCT_GUS_fwd | caaacaatgaAGTCTTCATCTTTTACACTTG |
| LK0321 | HR2_HCT_GUS_rev | ggtgacactatagagctagcGTACTCCTCATAAAGAATCTC |
| PCR screening of *PpHCT:uidA* transformants (direct PCR) | | |
| LK0396 | F1 | GCAATGGTAACGGAGACTTCA |
| LK0397 | R1 | TAACATACGGCGTGACATCG |
| LK0219 | F2 | GAAGCTCAGAGAATGCGATGG |
| LK0376 | R2 | CCGATCACCTGCGTCAAT |
| *PpHCT* disruption construct (GIBSON cloning) | | |
| LK0001 | HR1_HCT_KO_fwd | aatacgactcactatagggcgaattcCTTGTCTCATCCGCCAAA |
| LK0002 | HR1_HCT_KO_rev | gtcatagctgCAAAGTAGCACACGGACG |
| LK0003 | NPTII_fwd_HCT | tgctactttgCAGCTATGACCATGATTACGC |
| LK0004 | NPTII_rev_HCT | gatgaagactTTGGGTAACGCCAGGGTT |
| LK0005 | HR2_HCT_KO _fwd | cgttacccaaAGTCTTCATCTTTTACACTTGAACTG |
| LK0006 | HR2_HCT_KO _rev | tatttaggtgacactatagagaattcTGCTCTTCCAAGTTTGCATG |
| PCR screening of Δ*PpHCT* transformants (direct PCR) | | |
| LK0323 | F3 | CCGTTCAAGTCCACCATCACC |
| LK0227 | R3 | TGTCGTGCTCCACCATGTTG |
| LK0228 | F4 | AAATCCAGTGACCTGCAGGC |
| LK0324 | R4 | GAGCTCGCTAAAGGGTACCATAA |
| LK0229 | PpCYP98_F | GGCAGTCATGTGGGAGAACA |
| LK0230 | PpCYP98_R | ATGGCCCATTCCACCGAAAT |
| Molecular characterization of Δ*PpHCT* mutants (RT-PCR) | | |
| LK0273 | F5 | ATGGCCGCCGCAAGTCAAGTTC |
| LK0274 | R5 | TTAGAAGGATGCCACTAGTTTGG |
| LK0295 | L21_F | GGTTGGTCATGGGTTGCG |
| LK0296 | L21_R | GAGGTCAACTGTCTCGCC |
| qRT-PCR analysis | | |
| LK0340 | Pp4CL1_qF1 | CACCGGATTGCCGAAAGGTG |
| LK0341 | Pp4CL1_qR1 | ACCAACTCTCAACCCGCACA |
| LK0285 | PpHCT_qF1 | ACCACATGGACACATTTGCC |
| LK0286 | PpHCT_qR1 | ATTCATACGTGCGCTGCAAC |
| HR0577 | PpCYP98_qF1 | ATATGATCACGGCAGGCATG |
| HR0578 | PpCYP98_qR1 | TGAACATCCGGATTGCGAAC |

| | | |
|---|---|---|
| HR0849 | Pp3c19_1800_qF1 | ATGCTTGCATTGCAGTGCTG |
| HR0850 | Pp3c19_1800_qR1 | TTCAATGCGCGTGATAACCC |
| HR0855 | Pp3c27_3270_qF1 | AATTACGGTGCGCTTGATCC |
| HR0856 | Pp3c27_3270_qR1 | AAGCGCTTGATCAACGCATC |
| CDS cloning (Gateway) | | |
| HR1010 | Pp4CL1_attB1 | ggggacaagtttgtacaaaaaagcaggctTCATGTCTCCTAGTTTGCTCCCG |
| HR1011 | Pp4CL1_attB2 | ggggaccactttgtacaagaaagctgggtCCTATACTTTGTTTCTTAGATCCTTTCG |
| HR1030 | AtHCT_attB1 | ggggacaagtttgtacaaaaaagcaggctTCATGAAAATTAACATCAGAGATTCCACC |
| HR1031 | AtHCT_attB2 | ggggaccactttgtacaagaaagctgggtCTCATATCTCAAACAAAAACTTCTCA |
| HR0677 | PpHCT_AttB1 | ggggacaagtttgtacaaaaaagcaggctTCATGGCCGCCGCAAGTCAAG |
| HR0678 | PpHCT_AttB2 | ggggaccactttgtacaagaaagctgggtCTTAGAAGGATGCCACTAGTTTGG |
| HR1028 | MpHCT_attB1 | ggggacaagtttgtacaaaaaagcaggctTCATGGCAGGCTCGATGTGTTC |
| HR1029 | MpHCT_attB2 | ggggaccactttgtacaagaaagctgggtCCTACAGTTGGTGGATTAACTCGG |
| HR483 | PpCYP98_attB1 | ggggacaagtttgtacaaaaaagcaggctTCATGGCAGTCATGTGGGAG |
| HR484 | PpCYP98_attB2 | ggggaccactttgtacaagaaagctgggtCTCACGAAGGGGATGATCC |
| HR1012 | ATR1_attB1 | ggggacaagtttgtacaaaaaagcaggctTCATGACTTCTGCTTTGTATGCTTCC |
| HR1013 | ATR1_attB2 | ggggaccactttgtacaagaaagctgggtCTCACCAGACATCTCTGAGGT |
| *AtHCT* CRISPR | | |
| HR0969 | AtHCT_gRNA1_F | gattGCTCGGTGGCAGGCCGGACCA |
| HR0970 | AtHCT_gRNA1_R | aaacTGGTCCGGCCTGCCACCGAG**C** |
| HR0971 | AtHCT_CRISPR1_geno_F | CCTTCTGAGAGAGTTGGTCGAC |
| HR0972 | AtHCT_CRISPR1_geno_R | CTAGCTCGGAGGAGTGTTCG |

**Supplemental Table S5. List of multiple reaction monitoring methods used for metabolite analysis.**

| Molecule | Molecular weight | Ionization mode | Precursor ion | Collision energy | Product ion |
|---|---|---|---|---|---|
| *p*-coumaric acid | 164.2 | HESI- | 163.0 | 9 | 119.0 |
| caffeic acid | 180.2 | HESI+ | 179.0 | 18 | 135.1 |
| *p*-coumaroyl-threonate | 282.1 | HESI+ | 283.0 | 10 | 147.0 |
| *p*-coumaroyl-shikimate | 319.3 | HESI+ | 320.9 | 8 | 147.0 |
| | | HESI- | 318.9 | 8 | 163.1 |
| *p*-coumaroyl-quinate | 338.3 | HESI+ | 339.0 | 13 | 147.0 |
| *p*-coumaroyl-glucose | 326.3 | HESI- | 325.2 | 19 | 145.1 |
| caffeoyl-threonate | 298.1 | HESI+ | 299.0 | 38 | 89.0 |
| caffeoyl-shikimate | 336.1 | HESI+ | 337.4 | 13 | 163.0 |
| | | HESI- | 335.4 | 13 | 135.1 |
| caffeoyl-quinate | 354.3 | HESI+ | 354.9 | 11 | 163.0 |
| sinapoyl-malate | 340.3 | HESI+ | 341.2 | 10 | 207.0 |
| kaempferol 3-*O*-rhamnoside 7-*O*-rhamnoside | 578.5 | HESI+ | 579.0 | 27 | 287.1 |
| phenylalanine | 165.2 | HESI+ | 166.1 | 11 | 120.2 |
| malate | 134.1 | HESI- | 133.1 | 7 | 115.1 |
| threonate | 136.1 | HESI- | 135.1 | 17 | 60.4 |
| shikimate | 174.1 | HESI- | 173.1 | 10 | 93.2 |
| quinate | 192.2 | HESI- | 191.2 | 22 | 85.2 |

### Supplemental Table S6. Results of *t*-test from Figure 5H.

| Variable | P value | Mean of WT | Mean of ΔPpHCT | Difference | SE of difference | t ratio | df | Adjusted P Value |
|---|---|---|---|---|---|---|---|---|
| pC2T | 0.0036 | 1.000 | 2.635 | -1.635 | 0.3166 | 5.164 | 5.000 | 0.0078 |
| pC4T | 0.0169 | 0.9967 | 1.770 | -0.7733 | 0.2195 | 3.524 | 5.000 | 0.0169 |
| C2T | <0.0001 | 1.000 | 0.000 | 1.000 | 0.03381 | 29.57 | 5.000 | <0.0001 |
| C4T | 0.0026 | 0.9967 | 0.000 | 0.9967 | 0.1795 | 5.552 | 5.000 | 0.0078 |

### Supplemental Table S7. Results of *t*-test from Figure 5I.

| Variable | P value | Mean of WT | Mean of ΔPpHCT | Difference | SE of difference | t ratio | df | Adjusted P Value |
|---|---|---|---|---|---|---|---|---|
| pCA | 0.0012 | 1.003 | 2.900 | -1.897 | 0.2890 | 6.563 | 5.000 | 0.0012 |
| CA | 0.0001 | 1.000 | 0.000 | 1.000 | 0.09404 | 10.63 | 5.000 | 0.0003 |

### Supplemental Table S8. Results of *t*-test from Figure 5M.

| Variable | P value | Mean of WT | Mean of ΔPpHCT | Difference | SE of difference | t ratio | df | Adjusted P Value |
|---|---|---|---|---|---|---|---|---|
| p-coumarate | 0.0002 | 66.72 | 231.0 | -164.3 | 20.36 | 8.071 | 6.000 | 0.0023 |
| Caffeate | <0.0001 | 92.02 | 0.000 | 92.02 | 4.398 | 20.92 | 6.000 | <0.0001 |
| C16 | <0.0001 | 1326 | 424.9 | 901.5 | 31.59 | 28.54 | 6.000 | <0.0001 |
| C18 | <0.0001 | 239.1 | 46.50 | 192.6 | 9.190 | 20.96 | 6.000 | <0.0001 |
| C18:1 | <0.0001 | 219.0 | 55.12 | 163.9 | 9.615 | 17.04 | 6.000 | <0.0001 |
| C18:2 | 0.0003 | 95.93 | 54.75 | 41.18 | 5.512 | 7.472 | 6.000 | 0.0033 |
| C18:3 | 0.0182 | 50.68 | 32.51 | 18.17 | 5.645 | 3.219 | 6.000 | 0.0876 |
| C20 | <0.0001 | 199.9 | 41.81 | 158.1 | 6.397 | 24.71 | 6.000 | <0.0001 |
| C20:4 | 0.2891 | 83.85 | 92.14 | -8.290 | 7.131 | 1.163 | 6.000 | 0.6408 |
| C22 | <0.0001 | 321.1 | 88.58 | 232.5 | 9.303 | 24.99 | 6.000 | <0.0001 |
| C24 | <0.0001 | 246.8 | 89.95 | 156.9 | 3.861 | 40.63 | 6.000 | <0.0001 |
| C26 | <0.0001 | 98.41 | 14.54 | 83.88 | 3.523 | 23.81 | 6.000 | <0.0001 |
| 16-OH C16 | 0.0047 | 95.08 | 66.53 | 28.55 | 6.529 | 4.372 | 6.000 | 0.0279 |
| (9,10),16 di-OH C16 | 0.0025 | 789.8 | 112.0 | 677.8 | 136.1 | 4.982 | 6.000 | 0.0198 |
| 1,9,18-triOH C18 | 0.0009 | 32.63 | 13.52 | 19.10 | 3.147 | 6.071 | 6.000 | 0.0081 |
| 2-OH C20 | 0.0030 | 15.20 | 6.516 | 8.688 | 1.812 | 4.796 | 6.000 | 0.0209 |
| 2-OH C22 | 0.0884 | 40.68 | 30.27 | 10.41 | 5.122 | 2.032 | 6.000 | 0.3095 |
| 2-OH C24 | 0.8503 | 173.3 | 170.5 | 2.763 | 14.02 | 0.1971 | 6.000 | 0.8503 |
| 2-OH C24:1 | 0.4635 | 55.57 | 28.57 | 26.99 | 34.49 | 0.7828 | 6.000 | 0.7122 |
| 2-OH C26 | 0.0007 | 35.32 | 24.61 | 10.71 | 1.687 | 6.347 | 6.000 | 0.0071 |

**Supplemental Table S9. Results of *t*-test from Figure 6E.**

| Variable | P value | Mean of WT | Mean of $hct^{D7}$/*AtHCT* | Difference | SE of difference | t ratio | df | Adjusted P Value |
|---|---|---|---|---|---|---|---|---|
| pCG | <0.0001 | 1.000 | 81.80 | -80.80 | 2.937 | 27.51 | 6.000 | <0.0001 |
| pC5S | 0.0001 | 1.000 | 0.000 | 1.000 | 0.1151 | 8.687 | 6.000 | 0.0003 |
| C5S | <0.0001 | 1.000 | 0.000 | 1.000 | 0.08377 | 11.94 | 6.000 | <0.0001 |
| SM | <0.0001 | 1.000 | 0.1100 | 0.8900 | 0.04708 | 18.90 | 6.000 | <0.0001 |
| K3R7R | 0.0399 | 1.003 | 0.6950 | 0.3075 | 0.1176 | 2.615 | 6.000 | 0.0399 |
| Variable | P value | Mean of WT | Mean of $hct^{D7}$/*AtHCT* | Difference | SE of difference | t ratio | df | Adjusted P Value |
| pCG | 0.0002 | 1.000 | 48.24 | -47.24 | 5.813 | 8.128 | 6.000 | 0.0009 |
| pC5S | 0.0006 | 1.000 | 0.2525 | 0.7475 | 0.1154 | 6.480 | 6.000 | 0.0026 |
| C5S | 0.0013 | 1.000 | 0.3650 | 0.6350 | 0.1118 | 5.682 | 6.000 | 0.0038 |
| SM | 0.0018 | 1.000 | 0.7500 | 0.2500 | 0.04708 | 5.310 | 6.000 | 0.0038 |
| K3R7R | 0.4693 | 1.003 | 1.118 | -0.1150 | 0.1489 | 0.7722 | 6.000 | 0.4693 |
| Variable | P value | Mean of WT | Mean of $hct^{D7}$/*PpHCT* | Difference | SE of difference | t ratio | df | Adjusted P Value |
| pCG | 0.0030 | 1.000 | 25.54 | -24.54 | 5.107 | 4.805 | 6.000 | 0.0148 |
| pC5S | 0.0254 | 1.000 | 0.6225 | 0.3775 | 0.1276 | 2.958 | 6.000 | 0.0742 |
| C5S | 0.7096 | 1.000 | 0.9150 | 0.08500 | 0.2176 | 0.3906 | 6.000 | 0.8038 |
| SM | 0.0052 | 1.000 | 0.7650 | 0.2350 | 0.05500 | 4.273 | 6.000 | 0.0208 |
| K3R7R | 0.5570 | 1.003 | 0.9100 | 0.09250 | 0.1488 | 0.6217 | 6.000 | 0.8038 |
| Variable | P value | Mean of WT | Mean of $hct^{D7}$/*MpHCT* | Difference | SE of difference | t ratio | df | Adjusted P Value |
| pCG | 0.0002 | 1.000 | 16.14 | -15.14 | 1.811 | 8.358 | 6.000 | 0.0008 |
| pC5S | 0.0369 | 1.000 | 0.6250 | 0.3750 | 0.1403 | 2.673 | 6.000 | 0.1065 |
| C5S | >0.9999 | 1.000 | 1.000 | 0.000 | 0.1518 | 0.000 | 6.000 | >0.9999 |
| SM | 0.0029 | 1.000 | 0.7700 | 0.2300 | 0.04761 | 4.831 | 6.000 | 0.0116 |
| K3R7R | 0.9298 | 1.003 | 0.9900 | 0.01250 | 0.1360 | 0.09189 | 6.000 | 0.9951 |
